# spearfish: ultrafast Δ-mapping of long reads to redundant transcriptomes

**DOI:** 10.64898/2026.11.05.764698

**Authors:** Hesham Abdelgawad, Igor Martayan, Camille Marchet, Rob Patro

## Abstract

In a transcriptome, isoforms of a gene share most of their sequence, and gene families contain further near-copies. Yet most long-read mappers treat each isoform as a separate target, repeating seeding, chaining, and alignment across every shared exon.

We introduce `spearfish`, a long-read mapper built for this redundancy. Transcript quantification uses each read’s compatible transcripts, positions, and relative score differences rather than full alignments. `spearfish` therefore performs *delta mapping* (Δ-mapping): it computes each target’s alignment-score *difference* from the best candidate, skipping sequence shared by all candidates (which cancels out) and aligning differing regions once per distinct sequence rather than once per target. Seeds are *cores* from locally consistent parsing (LCP). Because each core depends only on local context, every isoform containing an exon has identical seeds across its interior. We collapse identical seeds within each gene into a *core graph*, a de Bruijn-like graph whose size is bounded by distinct exonic sequence plus a small term per splice junction, independent of isoform count. Over this graph, `spearfish` computes exact colinear chaining scores for every candidate transcript while sharing dynamic-programming state via a procedure provably equivalent to per-transcript chaining.

On 32 threads across RNA-seq datasets, `spearfish` maps reads 24–102× faster than `minimap2` in under a quarter of the memory, and 2.5–11.5× faster than `lr-kallisto`’s pseudoalignment. On simulated reads, it ranks the true transcript first more often than `minimap2` on HiFi reads and at least as often on ONT reads. With `oarfish`, abundance estimates match or improve on those from `minimap2` alignments on simulated reads (with or without primers and poly(A) tails) and seven SG-NEx spike-in runs, while quantifying 4.1–8.8× faster than `lr-kallisto`.

## 1 Introduction

Long-read RNA sequencing on Oxford Nanopore (ONT) and PacBio platforms can read RNA molecules end to end, making it possible in principle to assign each read to its transcript of origin. When an annotation is available, analyses such as transcript quantification typically begin by mapping reads to the annotated transcriptome. A long-read aligner such as `minimap2` [1] reports every transcript to which a read aligns nearly as well as its best hit, along with each alignment score. Quantifiers such as `oarfish` [2], transigner [3], NanoCount [4], and bambu [5] then resolve multi-mapping ambiguity with an expectation-maximization (EM) algorithm. Mapping dominates runtime: on a 26-million-read SG-NEx PCR-cDNA run [6], `minimap2` accounts for 99% of the CPU time of `minimap2` + `oarfish`.

Much of that work is redundant. GENCODE v50 [7] lists 670,670 human transcripts totaling 1.48 Gbp, yet a gene’s isoforms are assembled from a shared pool of exons. In our index, the 82.3 million seed occurrences across those transcripts collapse into 13.7 million distinct (seed, gene) pairs, and a seed match from a typical read occurs in ≈24 candidate transcripts. When a mapper treats transcripts as separate targets, as genome mappers treat chromosomes, it seeds, chains, and aligns the same exon once for every isoform containing it. Faster long-read genome mappers retain this repetition (Section 2).

Short-read mappers remove this redundancy by indexing a colored de Bruijn graph (where shared sequence is stored once and annotated with the set of transcripts, its *color*, containing it) and assigning reads to compatibility classes or tracking position sets jointly (Section 2). That works because short reads usually match a transcript nearly exactly or not at all. Long reads are harder: a read can span kilobases of shared sequence while isoform-distinguishing evidence is confined to one alternative splice site or a few bases at a transcript end. Sequencing errors (indels, homopolymers), untrimmed adapters, 5′ truncation, and chimeras easily obscure such differences, so positions and base-level scores matter.

What a quantifier actually uses from a mapper, however, is each read’s compatible transcripts, its position on each, and the relative score *differences* among candidates. Any score over sequence shared by all candidates cancels out. `spearfish` exploits this through *delta mapping* (Δ-mapping). Whereas pseudoalignment [8] determines compatibility and quasi-mapping [9] omits explicit scores, Δ-mapping reports every compatible transcript, the read’s position on it, and its score difference from the best candidate, omitting only the score common to all candidates. Shared sequence has the same score in every candidate and is skipped; differing sequence is evaluated once per distinct context. Three components are designed to realize this capability:

### 1. *Context-local seeds and the core graph* (Section 3, Section 4)

To collapse shared exons without losing error tolerance, we use locally consistent parsing (LCP) [10], which partitions a string into short overlapping *cores* determined only by their local neighborhood. Outside tandem repeats, an edit alters only nearby cores (about 3 per level on GENCODE). Sequencing errors therefore disrupt only adjacent seeds, and because a splice junction has the same local effect on the parse, every isoform containing an exon has identical seeds across its interior. We collapse identical seeds within each gene into a *core graph* storing each exon interior once plus a bounded number of nodes per junction (Theorem 2). Seeds whose neighborhood straddles an alternative splice boundary (the *junction halo*, Lemma 1) occur only in isoforms sharing that junction and distinguish isoforms differing by even a few bases. At matched density, an edit perturbs fewer LCP cores than minimizers (3.2 against 3.8), so fewer seeds are lost to errors (Section S1.1). Because LCP is hierarchical, `spearfish` seeds with level-3 cores and rescues noisy reads at level 2 (Section 5.2).

### 2. *Exact colinear chaining shared across isoforms* (Section 5.1)

Colinear chaining links seed matches (*anchors*) in transcript coordinates, where an exon absent from the read appears as a gap discrepancy in transcripts that include it. Because the chaining dynamic program (DP) depends on a transcript only through its relative anchor positions, isoforms with identical relative placements have identical DP values. We exploit this with a set-based refinement DP provably equivalent to per-transcript chaining (Theorem 3), and a faster prefix-sharing chainer built on the same principle.

### 3. *Base-level tie-breaking only where isoforms differ* (Section 5.3)

Transcripts with tied chaining scores are identical over most of the read. `spearfish` aligns only intervals between shared anchors where candidate sequences differ, evaluating each distinct sequence once. To prevent artifact bias, it soft-clips adapters, detects poly(A) tails, and aligns read ends uncompressed.

`spearfish` is written in Rust, runs standalone or inside `oarfish`, and is available under the BSD 3-clause license at https://github.com/COMBINE-lab/spearfish. On GENCODE v50, it maps reads 24–102× faster than `minimap2` in under 4 GB of memory with comparable accuracy (Section 6.1), and `oarfish` estimates from its mappings match or exceed those from `minimap2` alignments (Section 6.3).

## 2 Related work

### Fast short-read mapping

In a colored de Bruijn graph [11–14], every *k*-mer along a non-branching path (a *unitig*) occurs in the same set of transcripts. Short-read pseudoalignment [8] traces the unitigs matched by a read, skips uninformative regions, and intersects their transcript sets. Quasi-mapping [9, 12] adds read positions and colinearity, while selective alignment [15] aligns sequence between exact matches and caches identical subproblems. `lr-kallisto` [16] extends pseudoalignment to accurate long reads by increasing *k* (to 63) and falling back to the modal transcripts when strict intersection is empty. `spearfish` extends graph-based sharing to long-read chaining and alignment while keeping positions and score differences.

### Fast long-read mapping

Most work on faster long-read mapping targets `minimap2`’s seed-chain-extend pipeline on genomes [17]. Some tools keep `minimap2`’s output and speed up individual stages via SIMD kernels and a learned index (mm2-fast [18]), GPU chaining (mm2-gb [19]), or a Rust reimplementation (`rammap` [20], `oarfish`’s built-in mapper). Others change seeding: BLEND [21], minibwa [22], and mapquik [23], which indexes unique k-min-mers [24] (runs of *k* consecutive universe minimizers, *l*-mers whose hash falls below a threshold, occurring once in the reference) and chains them with a linear-time heuristic. mapquik maps human HiFi reads 37× faster than `minimap2`, but its accuracy drops sharply below 97% identity and reporting one alignment per read precludes transcriptome mapping. Others map raw nanopore signal [25] or long near-identical repeats [26].

Genome mappers assume one true locus per read, whereas a transcriptome mapper must report every compatible isoform and score near-ties. Dropping repeated k-min-mers, as mapquik does, discards seeds from shared exons. Meanwhile, `minimap2` accelerators retain those seeds but still chain and extend every candidate transcript separately. `spearfish` shares chaining and alignment across isoforms, works at ONT error rates, and runs banded alignment kernels only where candidates differ.

Other transcriptome-aware tools map to the genome first: uLTRA chains over annotated exons [27], IsoQuant [28] and bambu [5] assign spliced alignments to known or novel isoforms, and `bramble` [29] projects spliced alignments onto annotated transcripts for quantification.

## 3 Locally consistent parsing

To share work across isoforms under sequencing error, we need seeds placed identically wherever two sequences share a local neighborhood. Locally consistent parsing (LCP) [10, 30] partitions a string into short, overlapping *cores* using only local comparisons. At level 1, cores are placed at runs of identical characters and at local minima and maxima. Stretches without such landmarks are broken by deterministic coin tossing (DCT) [31], which iteratively maps adjacent symbols onto a smaller alphabet until landmarks appear. The parse is hierarchical: at level *k* + 1, the same rules are applied to the sequence of level-*k* core labels, so each level-(*k* + 1) core spans a short sequence of level-*k* cores (its *children*) and core length grows roughly geometrically. `spearfish` reimplements lcptools [32] and assigns each core a 64-bit label by hashing its bases (level 1) or its children’s labels and relative offsets (higher levels), so equal labels imply equal underlying sequence. Two properties of LCP are central:

### Determinism

A string’s parse depends only on the string itself, so away from its ends a query has the same parse as an identical substring of the reference.

### Local context

A level-*k* core spanning [*s, e*) is determined by its children and a few neighboring level-(*k −* 1) cores, so its label and boundaries depend only on the bases under them: its *extended window* [*s − λ, e* + *ρ*) with context radii *λ* and *ρ*. An edit at *p* alters only cores whose windows contain *p* (about 3.3 per level on GENCODE v50, where the 99th percentiles of *λ* and *ρ* at levels 2–7 are about 0.9 and 0.7 mean core lengths). Radii are bounded in cores rather than bases because tandem repeats can collapse into one long run core: in (*AC*)^*n*^, every level-1 core is *CAC*, forming one level-2 run core on which adjacent level-3 cores depend. Leaving long runs of equal symbols unparsed (a trade-off inherent to any bounded parse) bounds the radii in bases with unchanged accuracy (Section S1.1).

On raw RNA, homopolymer runs can be arbitrarily long (making even level-1 windows unbounded), and incorrect homopolymer run lengths are a major ONT error mode. *Homopolymer compression* (HPC), collapsing each run of identical bases to one, solves both: level-1 radii are at most 2 bases, and homopolymer-length errors vanish from the parse. Output positions are mapped back to raw coordinates.

LCP’s hierarchy involves a trade-off between specificity and error tolerance. Levels 1 and 2 are too short to be specific across a human transcriptome (level 1 has 466 distinct labels on GENCODE chr21 transcripts). Levels 4 and above are specific, but their wider windows more often contain sequencing errors and distinguish fewer isoform pairs (Section 4). Level 3 is the shortest level whose labels reliably identify a read’s gene (averaging ≈26 raw bases at ≈56 per kilobase), so `spearfish` indexes level-3 cores as its *seeds* (the ticks in Figure 1 a).

**Figure 1:**
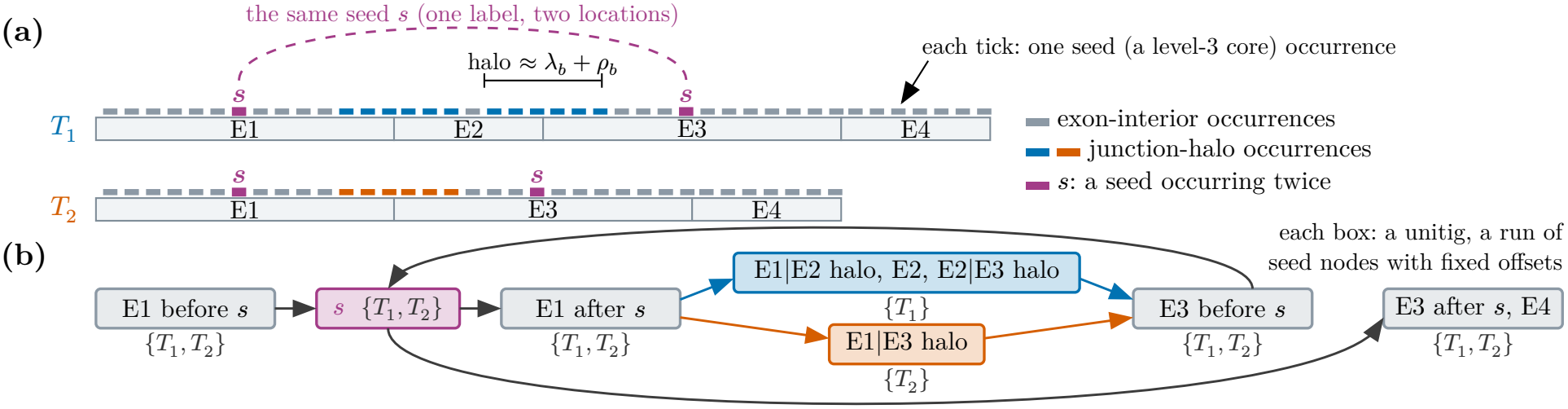
(a) Two isoforms of a gene in transcript coordinates. Interior seed occurrences whose extended windows lie inside a shared exon are invariant across isoforms (gray); halo occurrences near junctions may depend on the neighboring exon (blue or orange). The seed *s* (magenta) occurs twice in each isoform, in the E1 and E3 interiors. E4 follows E3 in both isoforms, so their junction has no halo and E3′s suffix and E4 form one unitig. (b) Equal (label, gene) pairs collapse to one node regardless of location. The compacted core graph therefore contains one shared *s* node, visited twice by each transcript, creating cycles. Transcript colors represent membership, not occurrence count.

## 4 The core graph

### 4.1 Construction

Once transcripts are parsed into level-3 seeds, shared exons produce identical seed runs across isoforms of a gene. Rather than indexing transcripts independently, we collapse repeated occurrences of each seed within a gene into a compacted *core graph* so each exon interior is stored once. Formally, let *T* = {*t*_1_, …, *t*_*m*_} be the transcripts and *b* = 3 the seed level. A *node* is a (seed label, gene) pair: all occurrences of a level-*b* label within one gene form one node, while occurrences in different genes remain distinct. Directed edges, labeled with the relative offset between seeds, connect consecutive nodes along any transcript’s parse. Maximal non-branching paths are compacted into *unitigs* (chains of seeds with fixed relative offsets), with transcript starts and ends treated as special predecessors and successors so every transcript enters and leaves unitigs only at endpoints. In this parse-induced precedence graph, consecutive seeds overlap by a variable amount stored on the edge. We call such graphs *de Bruijn-like* (for example, the syncmer graph of Durbin [33]). Like the transcriptome segments of Yanagi [34], each unitig *u* is traversed by a fixed set of transcripts, stored in an occurrence table occ(*u*) = [(*j*, off_*j*_(*u*))] listing each transcript traversing *u* and the offset of *u*’s first seed in *t*_*j*_ (with two entries if *t*_*j*_ traverses *u* twice in a tandem repeat). The *color* of *u* is the set of transcripts in occ(*u*). A seed at internal offset *δ* in *u* sits at

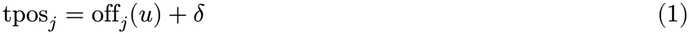

in every transcript *j* of *u*’s color, and chaining uses no other coordinate.

#### Why scope nodes by gene rather than collapsing labels globally?

About 14% of level-3 seeds occur in multiple genes. Global collapsing turns them into branch points, reducing unitig N50 from 36 seeds to 3, fragmenting chains, and roughly doubling mapping instructions (Table S6). With gene-scoped nodes, paralogs and repeats do not share unitigs, gene families need no special handling, and each gene is a natural chaining unit. Gene labels come from the annotation by default. Otherwise, `spearfish` clusters transcripts sharing seed runs, matching annotated genes and mapping accuracy on GENCODE v50.

The `spearfish` index stores the core graph, sorted 64-bit seed labels, occurrence and color tables, level-2 rescue tables (Section 5.2), and 2-bit transcript sequences in both HPC form (for internal alignment in parse coordinates) and uncompressed form (for read ends, Section 5.3). On GENCODE v50, it builds in 7 s on 32 threads and occupies 2.8 GB (1.5 GB rescue tables, 0.57 GB graph).

### 4.2 Invariant seeds and the junction halo

Two consequences of LCP’s local context determine both isoform resolution and graph size. Split a gene’s exons at every splice boundary used by any isoform into> non-overlapping *segments X* [35, 36], and let *J* be the set of distinct annotated junctions (pairs of consecutive segments). A segment boundary is *alternative* if isoforms containing the segment differ across it. Cores whose extended windows cross an alternative boundary form its *junction halo* (Figure 1).

#### Lemma 1 (halo).

*Let x* ∈ *X be shared by a set of isoforms I. A level-k core of x whose extended window lies inside x has the same label and the same offset relative to x in every isoform in I. A core whose extended window crosses an alternative boundary of x is shared only by isoforms that have the same sequence within the window beyond that boundary*.

Both parts follow because each core is a function of its extended window (Section S1.2): invariance breaks only within an extended window of an alternative boundary, holds across shared segment interiors, and breaks over a wider band at higher levels.

A halo seed occurs only in isoforms containing the adjacent junction, and its color is the set of those isoforms. A read from one isoform contains that isoform’s halo seeds and not the others’, so context-dependent seeds distinguish isoforms differing by only a few bases. Content-only samplers such as syncmers [37] depend solely on each *k*-mer’s bases and have no halo. At matched density, they distinguish 10–15 percentage points fewer isoform pairs differing by 1–30 nt than LCP or minimizers [38], whose window context yields a narrow halo. A wider halo is not better, though: higher-level cores span more context, so they more often contain a sequencing error and distinguish fewer pairs. `spearfish` therefore seeds with dense level-3 cores and rescues noisy reads using the LCP hierarchy (Section 5.2).

With shared interior cores and boundary-confined halos, graph size depends on distinct exonic sequence and junctions rather than total transcript length.

#### Theorem 2 (core-graph size).

*Let* 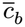 *be the minimum spacing of the gene’s level-b cores*, 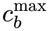 *their maximum length and λ*_*b*_, *ρ*_*b*_ *their largest window radii, and suppose every segment is longer than an extended window. The number of nodes in a gene’s core graph satisfies*

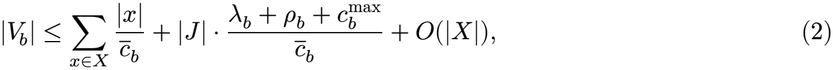

#### independent of the number of isoforms

An additional isoform built from existing exons contains no new exonic sequence, only new junctions. By Lemma 1, interior cores are counted once per segment. Every remaining core straddles a junction or transcript end and is determined by sequence within 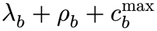 of that boundary, bounding labels per junction (Section S1.2). A per-transcript index without gene grouping instead grows with Σ_*j*_ |*t*_*j*_|. The bound applies to any local-window sampler: minimizer and syncmer windows have fixed width, while LCP’s vary yet remain explicit and measurable, as is the halo of Lemma 1.

## 5 Read mapping

`spearfish` maps each read in four stages (default thresholds in Section S3; pseudocode in Section S4): seeding and gene filtering, exact multi-transcript chaining, hierarchical level-2 rescue, and segmented base-level tie-breaking. It first homopolymer-compresses and parses the read, then looks up its level-3 seeds. Each match is an *anchor* (a pair of a query interval and a core-graph node). Anchors are grouped by gene, ignoring highly repetitive labels shared across too many genes.

Because a read typically comes from one gene or a small family, chaining every gene with an incidental seed match wastes time. `spearfish` therefore scores each gene greedily by summing anchor weights in query order (counting an overlapping anchor only past its predecessor), drops genes scoring well below the top gene, and retains the highest-scoring genes plus any close ties (up to a cap; Table S10). Keeping close ties is important in multi-copy families such as subtelomeric genes, where dozens of genes can have identical scores and a hard rank cutoff would exclude the true gene arbitrarily. Both read orientations are processed (sense strand only for direct RNA). Within each retained gene, consecutive read seeds matching consecutive positions of the same unitig with equal spacing in read and unitig are colinear with zero gap discrepancy in every transcript traversing that unitig, so `spearfish` merges them into one anchor before chaining.

### 5.1 Exact chaining in transcript coordinates, shared across isoforms

Unlike spliced genomic alignment, a read from an annotated transcript is expected to align contiguously, so consecutive anchors on the true isoform have nearly equal spacing in read and transcript coordinates (Δ*q* ≈ Δ*t*_*j*_). If a read comes from an isoform without exon *E* of candidate transcript *j*, a chain across that junction has discrepancy |Δ*q −* Δ*t*_*j*_| ≈ |*E*| in *j* and near zero in the true isoform. Together with halo-anchor colors, this discrepancy is the main chaining signal distinguishing isoforms. We therefore penalize |Δ*q −* Δ*t*_*j*_| linearly and omit the logarithmic gap term *β* log_2_(1 + max(Δ*q*, Δ*t*_*j*_)) [1] used for genomic introns (which are absent in transcript coordinates and whose inclusion lowers top-1 accuracy on simulated reads). Chains are local, so 5′-truncated reads are not penalized for missing sequence.

Formally, `spearfish` orders a gene’s anchors by query start. Anchor *i* has query interval [qs_*i*_, qe_*i*_), weight *w*_*i*_ (span capped at 100, slightly lowered for multi-gene seeds), length *L*_*i*_, and placements tpos_*j*_(*i*) in transcript *j* from Eq. 1 (one per unitig occurrence in *j*). A transition from anchor *i* to *i*′ > *i* in transcript *j* has query gap Δ*q* = qs_*i*′_ *−* qe_*i*_ and transcript gap Δ*t*_*j*_ = tpos_*j*_(*i*′) *−* (tpos_*j*_(*i*) + *L*_*i*_). The transition is valid if both gaps lie in [*−o*_max_, *g*_max_] and *i*′ *− i* ≤ *h*. Its gain is *w*_*i*′_ times the fraction of *i*′ not overlapping *i*, and its cost is

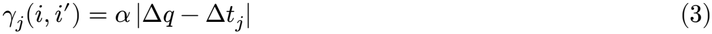

(*α* = 1.5, *h* = 50, *o*_max_ = 100, *g*_max_ = 5000). The best chain score *F*_*j*_ is the standard colinear chaining optimum [39, 40] in *j*’s coordinates. Because downstream quantifiers weight near-best candidates by relative score, we compute exact *F*_*j*_ for every valid candidate *j*, not just the top chain.

Chaining every isoform independently would repeat most of the dynamic programming. The DP for transcript *j* depends on *j* only through its anchor placements, and validity and cost (Eq. 3) depend only on placement *differences*. Two transcripts whose placement lists match up to a constant shift on anchors 1, …, *i* thus have identical DP values through anchor *i*: isoforms differing only upstream of the read (say, an alternative first exon) match across the entire read, and isoforms diverging inside the read match up to the first anchor past the divergence. On 10,000 simulated ONT reads at 95% accuracy, 36% of candidate transcripts share their entire shifted placement list with another candidate.

#### Prefix chainer

The prefix chainer uses this invariance directly: for each candidate, it takes the (anchor, placement) list from Eq. 1, subtracts the first placement from every entry, and sorts lists lexicographically. It chains the first transcript in full; for each later one, it reuses DP values along the longest anchor prefix shared with the preceding list and computes only the rest (Figure 2 a). By Proposition 4, this is exact. For example, ((*a*_1_, 3), (*a*_2_, 7), (*a*_3_, 10)) and ((*a*_1_, 4), (*a*_2_, 8), (*a*_3_, 11)) both normalize to ((*a*_1_, 0), (*a*_2_, 4), (*a*_3_, 7)), reusing all three DP values. A following list ((*a*_1_, 3), (*a*_2_, 7), (*a*_3_, 12)) matches only through *a*_2_, so its last entry is recomputed, while a transcript without *a*_1_ is chained from scratch. Sharing stops at the first difference, saving most when reads lie in shared exons.

**Figure 2:**
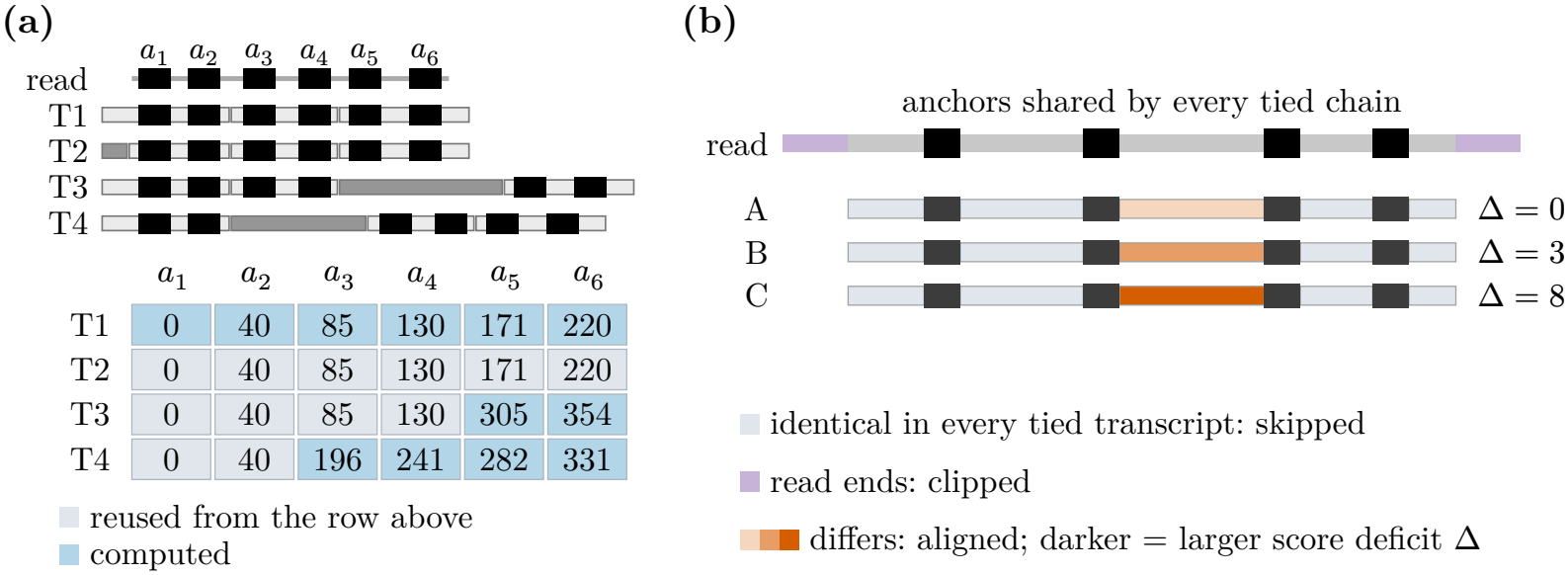
(a) A read’s six anchors placed in four transcripts (top): T2 differs from T1 only upstream of the read, while T3 and T4 have an extra exon (dark) after *a*_4_ and *a*_2_, respectively. The table lists the placements shifted so the first is 0, with rows sorted. Because chaining scores depend only on placement differences, the prefix chainer reuses the previous transcript’s DP values up to the first differing anchor (gray) and computes only the remainder (blue). (b) A separate example: for tied transcripts A–C, the tie-break aligns the read between shared anchors only where candidate sequences differ (orange, darker for a larger alignment-score deficit Δ relative to the best candidate); read ends are extended with soft-clipping, so adapter tails (purple) do not count.

#### Refinement DP

Even more state can be shared by grouping transcripts whenever their best scores coincide rather than only along common prefixes. In the *refinement DP*, each state is *σ* = (*i, D, f*), where *D* is a set of (transcript, placement) pairs of anchor *i* whose optimal chain ending at *i* has score *f* . Initially, each anchor *i* has one state containing all its placements with *f* = *w*_*i*_. To extend (*i, D, f*) to *i*′, we intersect *D* with *i*′’s color and partition by Δ*t*_*j*_. Within one unitig, offsets cancel so Δ*t*_*j*_ is identical for every *j*. Across unitigs, Δ*t*_*j*_ *−* (*δ*(*i*′) *− δ*_*i*_ *− L*_*i*_) = off_*j*_(*u*(*i*′)) *−* off_*j*_(*u*_*i*_) is the path length between the two unitigs in *j*, so each class consists of transcripts with equal path length between the anchors. (Paths differing in sequence but not length share a class, just as in per-transcript chaining; interior anchors or tie-breaking in Section 5.3 distinguish them.) Every member of a class has candidate score *f* + gain *− γ*. Keeping each placement’s maximum and grouping equal maxima forms the states of *i*′. Partitions depend only on the unitig pair and input class and are cached.

##### Theorem 3

(exact multi-transcript chaining). *With no beam (no cap on the number of states per anchor), for every anchor i, transcript j and placement p of i in j, exactly one state* (*i, D, f*) *of the refinement DP has* (*j, p*) ∈ *D, and f equals the per-transcript chaining optimum F*_*j*_(*i, p*) *with the same window, validity rule, gain and cost. Hence F*_*j*_ = max{*f* : (*i, D, f*), (*j*, ·) ∈ *D*} *for every j*.

The proof (Section S1.3) inducts on *i*, showing that candidate scores per placement match per-transcript chaining exactly. `spearfish` uses no beam cap because pruning low-scoring states would drop near-best candidates used by EM quantification.

Because equal prefixes imply equal scores, refinement-DP states are never finer than prefix-chainer classes, and merging diverging prefixes with equal scores yields fewer logical updates. Refinement-DP updates are set operations, however, whereas each prefix-chainer update runs a per-transcript DP loop scoring 16 predecessors per SIMD instruction. `spearfish` defaults to the prefix chainer and provides the refinement DP as an option. Both return exact optima for every candidate (Theorem 3, Proposition 4).

### 5.2 Rescue with the LCP hierarchy

Level-3 cores (≈26 raw bases) are rarely error-free below 85% identity, leaving the best chain covering under half the read. To recover such reads, `spearfish` re-parses the read at level 2 (≈11 raw bases). Single level-2 seeds are too common to index globally, yet within transcripts of candidate genes found at level 3 they are specific enough and remain intact more often at high error density. `spearfish` matches them locally within the window where the read can lie. For reads still without a half-length hit, the index tabulates runs of three consecutive level-2 seeds, which are specific on their own. When a best chain remains uncertain (few anchors, low coverage, or a close opposite-strand score; Section S4), `spearfish` re-chains it at level 2, including antisense transcripts. Below 80% identity on simulated reads, rescue raises true-transcript reporting from 60% to 92%, and disabling it lowers expressed-transcript Spearman correlation on 90% ONT reads from 0.847 to 0.810 (Table S6). On HiFi reads (≈0.1% error), rescue adds nothing and is off in the HiFi preset.

### 5.3 Base-level tie-breaking where isoforms differ

Chaining ranks transcripts by seed evidence, yet closely related isoforms (differing by a few bases, a short exon, or a shifted end) often tie or score nearly alike. Full-read alignment wastes work on shared sequence and adds error noise outside differing regions, so `spearfish` aligns selectively:

#### Alignment over distinct segments

The optimal chains of tied transcripts share common anchors. For each interval between consecutive shared anchors, `spearfish` extracts each tied transcript’s sequence. When all are identical (the usual case), every candidate has the same alignment cost and the interval is skipped. Where sequences differ, `spearfish` globally aligns one representative per distinct sequence with banded affine-gap alignment [41] and `minimap2`’s `map-ont` or `map-hifi` scores (Figure 2 b). Each candidate’s tie-break score is its best chain score minus its alignment-score deficit (Δ) relative to the best candidate. Whole-read tie-breaking executes 58% more instructions (Table S6), and segmented alignment is also more accurate because errors and divergence outside differing intervals affect all candidates equally and add noise.

#### Ties without a common anchor

Paralogs and repeats can tie with no anchor shared by all candidates. `spearfish` partitions them into anchor-sharing groups, runs segmented alignment within each group, and compares group representatives by the best contiguous run of banded alignments along each representative’s chain (with clipped ends as below). On HiFi reads, read-end bands are fixed on the anchor diagonal (low error).

#### End clipping

Because experimental reads often retain untrimmed adapters and primers, forcing end-to-end alignment penalizes true transcripts. `spearfish` aligns each read end (outside the outermost shared anchors) as a free-end extension with an X-drop and an end bonus for reaching the read end, soft-clipping unaligned flanks as in `minimap2`. When some candidates end before the read does, the end bonus applies only up to the shortest candidate’s end so stray flanking bases cannot inflate a longer isoform’s score.

#### Poly(A) tails

A poly(A) tail can spuriously align to A-rich sequence past a shorter isoform’s end. `spearfish` detects any A-rich tail near the read’s end past the last anchor, stops end alignment where the tail begins, and adds a small bonus to candidates whose annotated 3′ end (or an internal A-run) lies near that start (Table S10).

#### Read ends in uncompressed sequence

Template-switching cDNA protocols prepend `G` bases via the switch oligo. In HPC space, those `G`s are merged with a transcript’s leading `G`, so reads can align slightly better to an isoform annotated to start a few bases upstream. Except for direct RNA (no switch oligo), `spearfish` therefore aligns read ends against uncompressed transcript sequences stored in the index. On ONT cDNA, internal differing segments are also aligned uncompressed.

#### Read orientation

Antisense genes sharing sequence can give a read’s two orientations similar chain scores. When both orientations score close to one another, or when their best hits match identical sequence on opposite strands, `spearfish` compares them using clipped base-level alignment plus poly(A)-tail orientation evidence and reports both. When the margin remains small on ONT cDNA, it refines the comparison with banded whole-read local alignment, and for the quantification score (Section 5.4) it applies exact local alignment and a wider tail-flank tolerance to identical opposite-strand pairs (such as a short antisense isoform nested inside a longer one). Under the HiFi preset, if only one orientation has a poly(A) tail, `spearfish` maps that strand first and checks the opposite strand only if the best hit leaves a substantial part of the read uncovered. Exact margins, bonuses, and preset-specific cutoffs are listed in Section S3 and Algorithm S1.

### 5.4 Output and quantification

Because longer reads have more chances to accumulate spurious short chains by chance, `spearfish` leaves a read unmapped if its best chain score falls below a threshold that grows logarithmically with read length (Table S10). For mapped reads, it outputs every transcript within a generous score margin of the best, reporting score, strand, read and transcript intervals, and flags. Quantifiers use each candidate’s score difference from the best. Exact ties are ordered by a hash of read and transcript identifiers so ties are not systematically broken in index order. Reported intervals span the outermost anchors by default; with optional end extension (`--extend-ends`), full read coordinates are reported for downstream span filters or truncation models.

In the results below, `oarfish` estimates abundances via an in-process `spearfish` backend passing scored mappings directly to its EM in place of `minimap2` alignments. `oarfish` converts scores to conditional probabilities *P* (*r* | *t*_*j*_) *∝* exp((*s*_*j*_ *− s*_max_(*r*))/*D*) over hits within 95% of the read’s best (*D* = 2, default for both tools). Here *s*_*j*_ is `spearfish`’s *quantification score*: the mapping score, plus the orientation evidence above, plus a small *end fit* bonus (Table S10) when the read has no poly(A) tail, its aligned end reaches *t*_*j*_’s annotated 3′ end, and candidate 3′ ends differ. The end fit affects only EM weights, not reported mapping scores. Otherwise, the compared `oarfish` pipelines share the same quantification model.

#### RAD output

Mapping and quantification can also run separately: `spearfish` writes chunked binary RAD files [42] (one record per read with all its mappings, plus file-level tags for the quantifier). Because the layout is mapper-agnostic, other long-read mappers can write it too, and `oarfish` quantifies RAD faster than BAM.

## 6 Results

All benchmarks use the GENCODE v50 [7] human transcriptome, augmented with 333 SG-NEx synthetic spike-in sequences for the experimental runs. Simulated datasets were generated with PBSIM3 [43] across all transcripts at log-normal abundances: 1.97 M ONT reads at 95% mean accuracy (ERRHMM-ONT-HQ), 0.24 M ONT reads at 90% accuracy, and 1.94 M PacBio HiFi reads (multi-pass subreads processed with `ccs`). Experimental datasets comprise seven SG-NEx runs [6] spanning four protocols: ONT PCR-cDNA (HEYA8, H9), direct cDNA (HEYA8), direct RNA (HEYA8, H9), and PacBio (HEYA8, K562), evaluated against known concentrations of their synthetic spike-ins, SIRVs [44] and sequins [45].

We compare `spearfish` 0.3.1 (`ont`, `drna`, or `hifi` preset) against `minimap2` 2.31 (`-ax map-ont` or `map-hifi`, `-N 100`) and `lr-kallisto` (kallisto 0.52.0, bustools 0.45.1, *k* = 63 index). For `lr-kallisto`, each read’s mapping is its equivalence class from `kallisto bus --long --threshold 0.8`, numbered by read (`-n`) and exported with `bustools text -f`. All transcripts in the class count as tied for best, and reads without a class are unmapped. For quantification, `oarfish` 0.13.0 runs with default settings on `spearfish` mappings (`spearfish` + `oarfish`) and on `minimap2` alignments (`minimap2` + `oarfish`), while `lr-kallisto` runs `bustools count` and `quant-tcc --long` after `kallisto bus`. Before running `kallisto`, U bases are converted to T. All timings use 32 threads on an otherwise idle 128-core AMD EPYC 9555 server (16-thread results in Section S2.9).

### 6.1 Mapping accuracy

Because each simulated read’s transcript of origin is known, we evaluate mapping accuracy directly (Table 1). A read counts as mapped by `spearfish` if its best hit spans at least half the read. *Top-1* counts reads whose true transcript ties for the best score. Ties are expected because roughly a quarter of simulated reads originate from regions identical across multiple isoforms. *In set* checks whether the true transcript falls in the read’s *near-best set* (hits within 95% of the best score, or the equivalence class for `lr-kallisto`).

**Table 1:** Mapping accuracy on simulated reads. Mapped: best hit spans at least half the read (`lr-kallisto`: pseudoaligned). Top-1: the true transcript ties for the best score. In set: the true transcript is among the hits within 95% of the best score (`lr-kallisto`: its equivalence class); set size: their mean number. Gene: a best hit is in the true gene. ext.: spans extended to the read ends (`--extend-ends`). By read type in Table S1.

| **Reads** | **Mapper** | **Mapped** | **Top-1** | **In set** | **Set size** | **Gene** |
|---|---|---|---|---|---|---|
| ONT 95% | spearfish | **99.6%** | **99.2%** | **99.5%** | **5.0** | **99.5%** |
|  | minimap2 | **99.6%** | **99.2%** | **99.5%** | 7.3 | **99.5%** |
|  | lr-kallisto | 92.8% | n/a | 92.3% | 6.9 | 92.8% |
| HiFi | spearfish | **99.7%** | **99.6%** | 99.6% | 5.7 | 99.6% |
|  | spearfish, ext. | **99.7%** | **99.6%** | **99.7%** | 5.7 | **99.7%** |
|  | minimap2 | 98.5% | 98.3% | 98.4% | 6.1 | 98.5% |
|  | lr-kallisto | 96.8% | n/a | 96.6% | **4.3** | 96.8% |
| ONT 90% | spearfish | **98.2%** | **97.7%** | **98.0%** | **4.6** | **98.1%** |
|  | minimap2 | 98.1% | 97.6% | **98.0%** | 7.0 | **98.1%** |
|  | lr-kallisto | 58.5% | n/a | 58.1% | 20.1 | 58.5% |

On HiFi reads, `spearfish` outperforms `minimap2` on every measure, mapping 99.7% of reads versus 98.5% and ranking the true transcript first for 99.6% versus 98.3%. On ONT reads at both 95% and 90% accuracy, the two mappers stay within 0.1 percentage points of each other on every metric. Across all three simulated datasets, `spearfish` produces smaller near-best sets at virtually identical recall because its tie-break resolves isoforms whose `minimap2` scores lie within 5% of one another. Most remaining ambiguity occurs on 5′-truncated reads: on ONT 95% reads, full-length reads average 1.8 near-best transcripts compared to 7.9 for truncated reads (Table S1; breakdown by transcript category in Section S2.2). `lr-kallisto`’s class contains the true transcript for 99.2–99.8% of reads it pseudoaligns, but it pseudoaligns only 92.8% of ONT 95%, 58.5% of ONT 90%, and 96.8% of HiFi reads, far fewer than `spearfish` or `minimap2`.

On experimental reads, where ground truth is unknown, we compare the mappers on 200,000 random reads per SG-NEx run (Table S5). `spearfish` maps more cDNA reads than `minimap2` and 0.4–0.9 percentage points fewer direct RNA and PacBio reads. Among reads mapped by both tools, their near-best sets share at least one transcript for 99.0–99.7% of reads, and `spearfish`’s best hit falls in the gene of `minimap2`’s primary alignment for 98.3–99.6%. Consistent with simulation, `spearfish`’s near-best sets are roughly half as large on ONT reads and a fifth smaller on PacBio. `lr-kallisto` pseudoaligns 65–76% of these reads.

### 6.2 Speed and memory

`spearfish` maps reads 24–102× faster than `minimap2` (Table 2). The speedup is smallest on the PCR-cDNA runs, where rescue and tie-breaking are invoked most often because of adapter and chimeric sequences (stage-by-stage profiling in Section S2.8). `spearfish` is also 2.5–11.5× faster than `lr-kallisto`’s pseudoalignment step (`kallisto bus`) on every run. End to end from reads to transcript abundances, `spearfish` + `oarfish` is 11–65× faster than `minimap2` + `oarfish` and 4.1–8.8× faster than `lr-kallisto`, with similar mapping speedups at 16 threads (Section S2.9). Peak memory during mapping is 1.8–3.9 GB for `spearfish` (Table S7), compared to 13.5–20.2 GB for `minimap2` with its transcriptome index and 2.9–3.6 GB for `kallisto bus`.

**Table 2:** Wall-clock seconds on 32 threads. sf: `spearfish`; mm2: `minimap2` (index prebuilt); lrk: `lr-kallisto` (index prebuilt). Mapping: `spearfish`, `minimap2` and `lr-kallisto`’s pseudoalignment step (`kallisto bus`). Reads to abundances: ours, `oarfish` with the `spearfish` backend (including building the `spearfish` index, about 7 s); `minimap2` followed by `oarfish -a`; and `lr-kallisto`’s full pipeline. CPU times and memory are in Table S7.

|  |  | Mapping |  |  |  |  | Reads to abundances |  |  |  |
|---|---|---|---|---|---|---|---|---|---|---|
| Run | Reads | sf | mm2 | lrk | mm2 ÷ sf | lrk ÷ sf | ours | mm2 + oarfish | lrk | lrk ÷ ours |
| H9 direct RNA | 0.6 M | 1.6 | 100 | 18.6 | 62× | 11.5× | 9.2 | 102 | 76 | 8.3× |
| HEYA8 direct RNA | 1.0 M | 2.7 | 167 | 27 | 61× | 9.9× | 10.5 | 170 | 92 | 8.8× |
| HEYA8 direct cDNA | 8.5 M | 59 | 2,159 | 148 | 37× | 2.5× | 70 | 2,172 | 285 | 4.1× |
| HEYA8 PCR-cDNA | 25.9 M | 114 | 3,401 | 397 | 30× | 3.5× | 129 | 3,457 | 609 | 4.7× |
| H9 PCR-cDNA | 24.4 M | 131 | 3,117 | 377 | 24× | 2.9× | 147 | 3,154 | 613 | 4.2× |
| HEYA8 PacBio | 3.4 M | 12.8 | 1,308 | 58 | 102× | 4.5× | 20 | 1,312 | 85 | 4.2× |
| K562 PacBio | 2.2 M | 7.1 | 639 | 37 | 91× | 5.2× | 14.3 | 641 | 61 | 4.3× |
| Simulated ONT | 2.0 M | 5.8 | 436 | 36 | 75× | 6.2× | 13.2 | 439 | 75 | 5.7× |
| Simulated HiFi | 1.9 M | 4.0 | 329 | 18.1 | 82× | 4.5× | 11.7 | 331 | 51 | 4.4× |

### 6.3 Transcript quantification

We evaluate Δ-mapping end to end on transcript quantification, though we suspect other applications such as taxonomic assignment and metagenomic abundance estimation could also benefit from it. Here `oarfish`’s EM runs unchanged on `spearfish` mappings and on `minimap2` alignments, with `lr-kallisto` as a third pipeline.

On simulated reads, `spearfish` mappings yield more accurate estimates than `minimap2` alignments on every metric (Table 3), raising Spearman correlation by 0.03–0.05 and lowering mean absolute relative difference (MARD) by 0.02–0.03, especially on expressed transcripts where isoform resolution matters. Most of this gain is due to the end fit (Section 5.4), as clean PBSIM3 reads have no tail and end flush at their 3′ end. `lr-kallisto` is behind both `oarfish` pipelines on every metric, partly because it pseudoaligns only 92.8% of ONT reads (Table 1). With cDNA primers and poly(A) tails added (Section S2.3), the 3′-end signal comes from the tail instead and `spearfish` + `oarfish` is ahead by a slightly wider margin. We also evaluate fold-change accuracy across challenging transcript categories (Figure S1; Section S2.11).

**Table 3:** Simulated reads (PBSIM3). Spearman correlation and mean absolute relative difference (MARD) of estimated against true read counts, over all and over expressed transcripts; wall seconds from reads to abundances, 32 threads.

| **Reads** | **Pipeline** | **Spearman all / expr.** | **MARD all / expr.** | **Wall s** |
|---|---|---|---|---|
| ONT 95% | spearfish + oarfish | **0.913** / **0.876** | **0.114** / **0.159** | **13.2** |
|  | minimap2 + oarfish | 0.881 / 0.827 | 0.136 / 0.193 | 439 |
|  | lr-kallisto | 0.798 / 0.736 | 0.234 / 0.342 | 75 |
| HiFi | spearfish + oarfish | **0.915** / **0.877** | **0.111** / **0.157** | **11.7** |
|  | minimap2 + oarfish | 0.883 / 0.827 | 0.134 / 0.192 | 331 |
|  | lr-kallisto | 0.856 / 0.813 | 0.154 / 0.194 | 51 |

On the SIRV isoform sets, where isoforms of a gene must be told apart, `spearfish` + `oarfish` has the west MARD and detects the most isoforms in all seven SG-NEx runs (Table 4), including PacBio. On sequins, which span a range of abundances rather than focusing on isoform resolution, the two `oarfish` pipelines differ by at most 0.012 MARD (`spearfish` equal or lower on four of five runs). `lr-kallisto`’s lower sequin MARD on PacBio arises because it fails to pseudoalign most full-length reads of several long, over-represented sequins, coincidentally shifting estimates toward nominal concentrations (Section S2.6). Without span extension (Section 5.4), SIRV MARD rises on cDNA (0.432 vs. 0.422 on HEYA8 PCR-cDNA) because `oarfish`’s span filters apply to seed spans rather than read spans. Ablations appear in Section S2.5. On PCR-cDNA, estimates decline with isoform length relative to spike-ins. Adding a truncation model to `oarfish` (`--model-truncation`, Section S2.10) lowers SIRV MARD on all seven runs. The residual PCR-cDNA bias (absent in direct RNA) suggests amplification bias, a modeling rather than mapping issue.

**Table 4:** SG-NEx runs. sf: `spearfish` + `oarfish`; mm2: `minimap2` + `oarfish`; lrk: `lr-kallisto` (*k* = 63). MARD between estimated read counts and spike-in concentrations; detected: fraction of spike-in isoforms with at least one read. The isoform sets (SIRV-1, SIRV-4) contain several isoforms per gene; sequins span a range of abundances. ^†^Lower because `lr-kallisto` discards reads of long sequins over-represented in the libraries (Section S2.6).

|  |  | Isoform-set MARD |  |  | Isoforms detected |  |  | Sequin MARD |
|---|---|---|---|---|---|---|---|---|
| Run | Set | sf | mm2 | lrk | sf | mm2 | lrk | sf / mm2 / lrk |
| H9 direct RNA | SIRV-4 | **0.145** | 0.210 | 0.353 | **0.986** | 0.899 | 0.884 | n/a |
| HEYA8 direct RNA | SIRV-1 E2 | **0.240** | 0.282 | 0.355 | **0.971** | 0.884 | 0.899 | **0.829** / 0.833 / 0.870 |
| HEYA8 direct cDNA | SIRV-1 E2 | **0.293** | 0.325 | 0.332 | **0.986** | 0.913 | 0.928 | **0.372** / 0.384 / 0.381 |
| HEYA8 PCR-cDNA | SIRV-1 E2 | **0.422** | 0.453 | 0.462 | **0.986** | 0.913 | 0.928 | 0.386 / **0.382** / 0.396 |
| H9 PCR-cDNA | SIRV-4 | **0.382** | 0.405 | 0.406 | **1.000** | 0.913 | 0.928 | n/a |
| HEYA8 PacBio | SIRV-4 | **0.771** | 0.783 | 0.787 | **0.928** | 0.899 | 0.884 | 0.772 / 0.773 / **0.754**<sup>†</sup> |
| K562 PacBio | SIRV-4 | **0.768** | 0.781 | 0.792 | **0.971** | 0.913 | 0.913 | 0.746 / 0.747 / **0.730**<sup>†</sup> |

## 7 Discussion

We introduced Δ-mapping and implemented it in `spearfish`, reporting positions and relative score differences while evaluating shared sequence once: the core graph stores each exon interior once, the chainer evaluates each distinct anchor-placement list once, and the tie-break aligns only where tied isoforms differ. `spearfish` maps 24–102× faster than `minimap2` with comparable accuracy, and `oarfish` estimates from its mappings match or improve on `minimap2` alignments.

We chose LCP seeds because they work well in practice, yet Δ-mapping is not tied to them: at matched density, level-3 cores and minimizers have the same graph-size bound and isoform resolution, making other seeding schemes promising future work. `spearfish` could also extend to related genome collections (e.g. metagenomics) or spliced genome alignment.

`spearfish`’s main limitations are on very low-identity reads (below 80% identity, reads are still mapped less often than with `minimap2`) and scope: it outputs compatibility sets, placements, and relative scores rather than full alignments, so it is a front end where Δ-mapping suffices rather than a general-purpose aligner. For those tasks, it matches or exceeds full-aligner accuracy in a fraction of the time and memory.

## Declarations

RP is a co-founder of Ocean Genomics Inc.

## Acknowledgements

This work was supported by the US National Institutes of Health (R01HG009937) and by grants 2022-311195 and 2024-342821 from the Chan Zuckerberg Initiative DAF, an advised fund of the Chan Zuckerberg Initiative Foundation.

## References

1. Li, H.: Minimap2: pairwise alignment for nucleotide sequences. Bioinformatics. 34, 3094–3100 (2018). 10.1093/bioinformatics/bty191.

2. Zare Jousheghani, Z., Singh, N.P., Patro, R.: Oarfish: Enhanced probabilistic modeling leads to improved accuracy in long read transcriptome quantification. Bioinformatics. 41, i304–i313 (2025). 10.1093/bioinformatics/btaf240.

3. Ji, H.J., Pertea, M.: Enhancing transcriptome expression quantification through accurate assignment of long RNA sequencing reads with TranSigner. Genome Biology. 26, 257 (2025). 10.1186/s13059-025-03723-2.

4. Gleeson, J., Leger, A., Prawer, Y.D.J., Lane, T.A., Harrison, P.J., Haerty, W., Clark, M.B.: Accurate expression quantification from nanopore direct RNA sequencing with NanoCount. Nucleic Acids Research. 50, e19 (2022). 10.1093/nar/gkab1129.

5. Chen, Y., Sim, A., Wan, Y.K., Yeo, K., Lee, J.J.X., Ling, M.H., Love, M.I., Göke, J.: Context-aware transcript quantification from long-read RNA-seq data with Bambu. Nature Methods. 20, 1187–1195 (2023). 10.1038/s41592-023-01908-w.

6. Chen, Y., Davidson, N.M., Wan, Y.K., Yao, F., Su, Y., Gamaarachchi, H., Sim, A., Patel, H., Low, H.M., Hendra, C., Wratten, L., Hakkaart, C., Sawyer, C., Iakovleva, V., Lee, P.L., Xin, L., Ng, H.E.V., Loo, J.M., Ong, X., Ng, H.Q.A., Wang, J., Koh, W.Q.C., Poon, S.Y.P., Stanojevic, D., Tran, H.-D., Lim, K.H.E., Toh, S.Y., Ewels, P.A., Ng, H.-H., Iyer, N.G., Thiery, A., Chng, W.J., Chen, L., DasGupta, R., Sikic, M., Chan, Y.-S., Tan, B.O.P., Wan, Y., Tam, W.L., Yu, Q., Khor, C.C., Wüstefeld, T., Lezhava, A., Pratanwanich, P.N., Love, M.I., Goh, W.S.S., Ng, S.B., Oshlack, A., SG-NEx consortium, Göke, J.: A systematic benchmark of Nanopore long-read RNA sequencing for transcript-level analysis in human cell lines. Nature Methods. 22, 801–812 (2025). 10.1038/s41592-025-02623-4.

7. Frankish, A., Carbonell-Sala, S., Diekhans, M., Jungreis, I., Loveland, J.E., Mudge, J.M., Sisu, C., Wright, J.C., Arnan, C., Barnes, I., Banerjee, A., Bennett, R., Berry, A., Bignell, A., Boix, C., Calvet, F., Cerdán-Vélez, D., Cunningham, F., Davidson, C., Donaldson, S., Dursun, C., Fatima, R., Giorgetti, S., Giron, C.G., Gonzalez, J.M., Hardy, M., Harrison, P.W., Hourlier, T., Hollis, Z., Hunt, T., James, B., Jiang, Y., Johnson, R., Kay, M., Lagarde, J., Martin, F.J., Gómez, L.M., Nair, S., Ni, P., Pozo, F., Ramalingam, V., Ruffier, M., Schmitt, B.M., Schreiber, J.M., Steed, E., Suner, M.-M., Sumathipala, D., Sycheva, I., Uszczynska-Ratajczak, B., Wass, E., Yang, Y.T., Yates, A., Zafrulla, Z., Choudhary, J.S., Gerstein, M., Guigo, R., Hubbard, T.J.P., Kellis, M., Kundaje, A., Paten, B., Tress, M.L., Flicek, P.: GENCODE: reference annotation for the human and mouse genomes in 2023. Nucleic Acids Research. 51, D942–D949 (2023). 10.1093/nar/gkac1071.

8. Bray, N.L., Pimentel, H., Melsted, P., Pachter, L.: Near-optimal probabilistic RNA-seq quantification. Nature Biotechnology. 34, 525–527 (2016). 10.1038/nbt.3519.

9. Srivastava, A., Sarkar, H., Gupta, N., Patro, R.: RapMap: a rapid, sensitive and accurate tool for mapping RNA-seq reads to transcriptomes. Bioinformatics. 32, i192–i200 (2016). 10.1093/bioinformatics/btw277.

10. Sahinalp, S.C., Vishkin, U.: Efficient approximate and dynamic matching of patterns using a labeling paradigm. In: Proceedings of the 37th Annual Symposium on Foundations of Computer Science (FOCS). pp. 320–328 (1996). 10.1109/sfcs.1996.548491.

11. Liu, B., Guo, H., Brudno, M., Wang, Y.: deBGA: read alignment with de Bruijn graph-based seed and extension. Bioinformatics. 32, 3224–3232 (2016). 10.1093/bioinformatics/btw371.

12. Almodaresi, F., Sarkar, H., Srivastava, A., Patro, R.: A space and time-efficient index for the compacted colored de Bruijn graph. Bioinformatics. 34, i169–i177 (2018). 10.1093/bioinformatics/bty292.

13. Khan, J., Patro, R.: Cuttlefish: fast, parallel and low-memory compaction of de Bruijn graphs from large-scale genome collections. Bioinformatics. 37, i177–i186 (2021). 10.1093/bioinformatics/btab309.

14. Fan, J., Khan, J., Singh, N.P., Pibiri, G.E., Patro, R.: Fulgor: a fast and compact k-mer index for large-scale matching and color queries. Algorithms for Molecular Biology. 19, 3 (2024). 10.1186/s13015-024-00251-9.

15. Srivastava, A., Malik, L., Sarkar, H., Zakeri, M., Almodaresi, F., Soneson, C., Love, M.I., Kingsford, C., Patro, R.: Alignment and mapping methodology influence transcript abundance estimation. Genome Biology. 21, 239 (2020). 10.1186/s13059-020-02151-8.

16. Loving, R.K., Sullivan, D.K., Reese, F., Rebboah, E., Sakr, J., Rezaie, N., Liang, H.Y., Filimban, G., Kawauchi, S., Booeshaghi, A.S., Melsted, P., Oakes, C., Trout, D., Williams, B.A., MacGregor, G.R., Wold, B.J., Mortazavi, A., Pachter, L.: Long-read sequencing transcriptome quantification with lr-kallisto. PLOS Computational Biology. 21, e1013692 (2025). 10.1371/journal.pcbi.1013692.

17. Sahlin, K., Baudeau, T., Cazaux, B., Marchet, C.: A survey of mapping algorithms in the long-reads era. Genome Biology. 24, 133 (2023). 10.1186/s13059-023-02972-3.

18. Kalikar, S., Jain, C., Vasimuddin, M., Misra, S.: Accelerating minimap2 for long-read sequencing applications on modern CPUs. Nature Computational Science. 2, 78–83 (2022). 10.1038/s43588-022-00201-8.

19. Dong, J., Liu, X., Sadasivan, H., Sitaraman, S., Narayanasamy, S.: mm2-gb: GPU accelerated minimap2 for long read DNA mapping. In: Proceedings of the 15th ACM International Conference on Bioinformatics, Computational Biology and Health Informatics (BCB). pp. 1–9 (2024). 10.1145/3698587.3701366.

20. Wang, J.R., Li, H.: Memory-safe high-performance sequence mapping with rammap. bioRxiv. (2026). 10.64898/2026.05.26.726289.

21. Firtina, C., Park, J., Alser, M., Kim, J.S., Cali, D.S., Shahroodi, T., Ghiasi, N.M., Singh, G., Kanellopoulos, K., Alkan, C., Mutlu, O.: BLEND: a fast, memory-efficient and accurate mechanism to find fuzzy seed matches in genome analysis. NAR Genomics and Bioinformatics. 5, lqad004 (2023). 10.1093/nargab/lqad004.

22. Li, H., Homer, N.: Fast genomic read alignment with minibwa. arXiv. 2606.15357 (2026). 10.48550/arxiv.2606.15357.

23. Ekim, B., Sahlin, K., Medvedev, P., Berger, B., Chikhi, R.: Efficient mapping of accurate long reads in minimizer space with mapquik. Genome Research. 33, 1188–1197 (2023). 10.1101/gr.277679.123.

24. Ekim, B., Berger, B., Chikhi, R.: Minimizer-space de Bruijn graphs: Whole-genome assembly of long reads in minutes on a personal computer. Cell Systems. 12, 958–968.e6 (2021). 10.1016/j.cels.2021.08.009.

25. Firtina, C., Soysal, M., Lindegger, J., Mutlu, O.: RawHash2: mapping raw nanopore signals using hash-based seeding and adaptive quantization. Bioinformatics. 40, btae478 (2024). 10.1093/bioinformatics/btae478.

26. Jain, C., Rhie, A., Hansen, N.F., Koren, S., Phillippy, A.M.: Long-read mapping to repetitive reference sequences using Winnowmap2. Nature Methods. 19, 705–710 (2022). 10.1038/s41592-022-01457-8.

27. Sahlin, K., Mäkinen, V.: Accurate spliced alignment of long RNA sequencing reads. Bioinformatics. 37, 4643–4651 (2021). 10.1093/bioinformatics/btab540.

28. Prjibelski, A.D., Mikheenko, A., Joglekar, A., Smetanin, A., Jarroux, J., Lapidus, A.L., Tilgner, H.U.: Accurate isoform discovery with IsoQuant using long reads. Nature Biotechnology. 41, 915–918 (2023). 10.1038/s41587-022-01565-y.

29. Rudnick, Z., Varabyou, A., Patro, R., Pertea, M.: Bramble: projection of spliced genomic alignments into transcriptomic space for improved transcript quantification. bioRxiv. (2026). 10.64898/2026.09.09.750464.

30. Sahinalp, S.C., Vishkin, U.: Symmetry breaking for suffix tree construction. In: Proceedings of the 26th Annual ACM Symposium on Theory of Computing (STOC). pp. 300–309 (1994). 10.1145/195058.195164.

31. Cole, R., Vishkin, U.: Deterministic coin tossing with applications to optimal parallel list ranking. Information and Control. 70, 32–53 (1986). 10.1016/s0019-9958(86)80023-7.

32. Ashyralyyev, A., Bingöl, Z., Öz, B.F., Zhu, K., Malikic, S., Vishkin, U., Sahinalp, S.C., Alkan, C.: LCPan: efficient variation graph construction using locally consistent parsing. Genome Biology. 27, 197 (2026). 10.1186/s13059-026-04088-w.

33. Durbin, R.: A Run-Length-Compressed Skiplist Data Structure for Dynamic GBWTs Supports Time-and Space-Efficient Pangenome Operations over Syncmers. Journal of Computational Biology. 15578666261495122 (2026). 10.1177/15578666261495122.

34. Gunady, M.K., Mount, S.M., Corrada Bravo, H.: Yanagi: Fast and interpretable segment-based alternative splicing and gene expression analysis. BMC Bioinformatics. 20, 421 (2019). 10.1186/s12859-019-2947-6.

35. Feng, J., Li, W., Jiang, T.: Inference of Isoforms from Short Sequence Reads. In: Research in Computational Molecular Biology (RECOMB 2010). pp. 138–157. Springer (2010). 10.1007/978-3-642-12683-3_10.

36. Feng, J., Li, W., Jiang, T.: Inference of Isoforms from Short Sequence Reads. Journal of Computational Biology. 18, 305–321 (2011). 10.1089/cmb.2010.0243.

37. Edgar, R.: Syncmers are more sensitive than minimizers for selecting conserved k-mers in biological sequences. PeerJ. 9, e10805 (2021). 10.7717/peerj.10805.

38. Roberts, M., Hayes, W., Hunt, B.R., Mount, S.M., Yorke, J.A.: Reducing storage requirements for biological sequence comparison. Bioinformatics. 20, 3363–3369 (2004). 10.1093/bioinformatics/bth408.

39. Abouelhoda, M.I., Ohlebusch, E.: Chaining algorithms for multiple genome comparison. Journal of Discrete Algorithms. 3, 321–341 (2005). 10.1016/j.jda.2004.08.011.

40. Jain, C., Gibney, D., Thankachan, S.V.: Co-linear chaining with overlaps and gap costs. In: Research in Computational Molecular Biology (RECOMB). pp. 246–262 (2022). 10.1007/978-3-031-04749-7_15.

41. Gotoh, O.: An improved algorithm for matching biological sequences. Journal of Molecular Biology. 162, 705–708 (1982). 10.1016/0022-2836(82)90398-9.

42. He, D., Zakeri, M., Sarkar, H., Soneson, C., Srivastava, A., Patro, R.: Alevin-fry unlocks rapid, accurate and memory-frugal quantification of single-cell RNA-seq data. Nature Methods. 19, 316–322 (2022). 10.1038/s41592-022-01408-3.

43. Ono, Y., Hamada, M., Asai, K.: PBSIM3: a simulator for all types of PacBio and ONT long reads. NAR Genomics and Bioinformatics. 4, lqac092 (2022). 10.1093/nargab/lqac092.

44. Paul, L., Kubala, P., Horner, G., Ante, M., Holländer, I., Seitz, A., Reda, T.: SIRVs: Spike-In RNA Variants as external isoform controls in RNA-sequencing. bioRxiv. 080747 (2016). 10.1101/080747.

45. Hardwick, S.A., Chen, W.Y., Wong, T., Deveson, I.W., Blackburn, J., Andersen, S.B., Nielsen, L.K., Mattick, J.S., Mercer, T.R.: Spliced synthetic genes as internal controls in RNA sequencing experiments. Nature Methods. 13, 792–798 (2016). 10.1038/nmeth.3958.

46. Myers, G.: A fast bit-vector algorithm for approximate string matching based on dynamic programming. Journal of the ACM. 46, 395–415 (1999). 10.1145/316542.316550.

47. Ukkonen, E.: Algorithms for approximate string matching. Information and Control. 64, 100–118 (1985). 10.1016/s0019-9958(85)80046-2.

