## Supplementary Information for "spearfish: ultrafast Δ-mapping of long reads to redundant transcriptomes"

#### S1 Proofs

##### S1.1 Extended windows

At level 1 after HPC, whether a core starts at position $i$ is determined by at most $S[i-1, i+4)$, and a run core contains its entire run, so a level-1 core depends on at most two bases on either side. At level $k > 1$, the symbol of a level-$(k-1)$ core is its DCT value (a function of its own bits and those of its left neighbor), and each level-$k$ core is determined by the same parsing rules from its children's symbols and a few level-$(k-1)$ cores on either side. The *extended window* $W(c) = [s - \lambda, e + \rho)$ of a level-1 core $c = [s, e)$ is the set of bases that determine it under the parsing rules; for a level-$k$ core, $W(c)$ is the union of the extended windows of all level-$(k-1)$ cores on which $c$ depends via the parsing rules and DCT. By induction on $k$, any string matching $S$ on $W(c)$ (away from its ends) has the same level-$(k-1)$ cores there and hence also $c$, with the same label at the same offset. This invariance is the property used to prove Lemma 1.

The extended window is bounded in numbers of cores, not bases. Consider $S = P(AC)^n Q$ with $P =$ `GTACATCGCTACGTCACTGCTGAT` and $Q =$ `GTGACTGATAGCGATATAGCGTAC`, both unchanged under HPC. Every level-1 core inside the repeat is $CAC$, so the level-2 input is a run of identical symbols. Under the run rule, that run forms a single level-2 core spanning the entire repeat, just as a homopolymer forms one level-1 core without HPC. Because the level-3 core immediately to the right of the repeat depends on this level-2 core as a neighbor, an edit just to the left of the repeat alters it. For $n = 10, 20, 40, 80$, the largest $\lambda_3$ over all single edits is $19, 39, 79, 159 = 2n - 1$ bases, while the level-1 and level-2 radii remain at most 2 and 4. With HPC, radii at levels 1 and 2 are bounded by a constant, whereas from level 3 onward they scale with the longest level-2 core. Any tandem repeat whose level-1 parse repeats a single core yields such a long level-2 core.

In real transcripts, windows become long in bases only adjacent to such repeats. Across 30,000 GENCODE v50 transcripts subjected to 9 random edits each, every changed core at levels 2–5 lies within 5 cores of a changed core at the level below, and over 90% overlap one directly. Across all of GENCODE v50, only 639 of 194 M level-2 cores span 100 bases or more (125 span at least 100 compressed bases; the longest, 177 compressed bases, occurs in an *RPGR* isoform). On the 655,487 transcripts of at least 200 compressed bases with 9 random edits each, the maximum $\lambda_3$ and $\rho_3$ are 116 and 65 compressed bases, compared to a mean level-3 core length of 18.6. At matched seed density, fewer seeds change per edit than with fixed-window minimizers: on the same transcripts, a substitution alters 3.22 level-3 cores (56 per raw kb) versus 3.76 minimizers with $k = 19, w = 23$ (58 per raw kb), and on chr21 transcripts with 6% simulated ONT errors, 41% of level-3 cores remain unchanged versus 38% of minimizers.

*Tandem repeats.* Every locally consistent parse is subject to this trade-off. If whether a phrase boundary occurs at position $i$ depends only on $S[i - R, i + R)$, then inside an exact tandem repeat of period $P$ longer than $2R$, those local windows, and therefore the boundaries, repeat with period $P$. Any phrases there must span at most one period plus their overlap, or there are none. A parse with both bounded context and bounded phrase lengths must therefore emit either short phrases or no phrases inside short-period repeats. Standard LCP's run rule instead forms a single long core, giving up the base-level context bound.

Capping the run length restores the bound. If any run of more than $m$ equal symbols (at any level) forms no core, with parsing resuming on either side as at a sequence boundary, then every run core has at most $m+2$ children, and the gap between consecutive landmarks is less than roughly twice the DCT alphabet size. By induction, every core then has a bounded number of children and an extended window bounded in bases by a constant depending only on $m$ and the level. Although the theoretical constant from this argument is large, empirical radii are small: with $m = 4$ and HPC, $\lambda_3 = 3$ on $P(AC)^n Q$ for every $n \geq 20$. On the 655,487 transcripts with 3 random edits of each type, maximum $\lambda_3$ and $\rho_3$ drop from 116 and 65 to 40 and 33 compressed bases, their 99th percentiles shift by at most one base, the longest level-3 core drops from 203 to 52 compressed bases, no core has more than 8 children, and only 0.69% of compressed bases between a transcript's first and last level-3 core are uncovered. Mapping and quantification accuracy are unchanged (Table S6), and 32-thread wall time across the three simulated sets stays within 2%. Because the worst-case base-level radius is mainly of theoretical rather than practical consequence on these data, we retain the standard LCP rules and leave modified parsing rules to future work.

##### S1.2 Lemma 1 and Theorem 2

*Proof.* (Lemma 1) Let $c$ be a level-$k$ core of segment $x$ whose extended window $W$ lies inside $x$. In any isoform containing $x$, the bases of $W$ are the bases of $x$ at the same offsets, and $c$ is a function of these bases (Section S1.1), so the parse has a core with the same boundaries and label at the same offset relative to $x$. If instead $W$ crosses an alternative boundary of $x$, the label and boundaries depend on the bases beyond the boundary, which differ between isoforms with different neighbors, so $c$ is shared only by isoforms that have the same sequence within $W$.

(Theorem 2) Partition the cores of the gene's transcripts by their extended windows. Cores whose window lies inside a segment are, by Lemma 1, determined by the segment and an offset. Consecutive cores start at least $\overline{c}_b$ apart, so there are at most $|x| / \overline{c}_b$ distinct nodes from segment $x$. Every other core has a window that contains a segment boundary. A core has a boundary in its window only if its span starts within $\lambda_b + \rho_b + c_b^{\max}$ before or after that boundary. Since every segment is longer than a window, its window contains exactly one boundary, which is either a junction $(x, y) \in J$ or one of the $O(|X|)$ transcript ends, and the window lies inside $xy$. The core is then determined by the junction and an offset, and offsets within that band take at most $(\lambda_b + \rho_b + c_b^{\max})/\overline{c}_b$ values. Summing over segments, junctions, and ends gives the bound, in which the number of isoforms does not appear. The parameters $\lambda_b$, $\rho_b$, and $c_b^{\max}$ are maxima over the gene's cores, so with an additional isoform they can change only through the cores of its new junctions. □

##### S1.3 Theorem 3

Fix a candidate gene with anchors $1, \ldots, n$ in query order. For transcript $j$ and placement $p$ of anchor $i$, the per-transcript DP is

$$F_j(i,p) = \max\left\{ w_i, \max_{(i',p')} F_j(i',p') + g(i',i) - \gamma_j(i',i) \right\}, \tag{S1}$$

taken over valid predecessors: anchors $i'$ with $i - h \leq i' < i$ and placements $p'$ of $i'$ in $j$ for which the transition is valid. The gain $g(i', i)$ depends only on query intervals.

*Proof.* By induction on $i$, we show that the states of $i$ partition the pairs $(j, p)$ and that the state containing $(j, p)$ has $f = F_j(i, p)$. For $i = 1$, the single state has $f = w_1 = F_j(1, p)$. Assume the claim holds for all anchors before $i$, and fix $(j, p)$. Every candidate value for $(j, p)$ in the refinement DP comes from a state $(i', D, f)$ with $i - h \leq i' < i$ and a class $C$ of its refinement containing a pair $(j, p')$ paired with placement $p$ of $i$ in $j$. Within $C$, all pairs have the same $\Delta t$, hence the same validity and cost, so $f + g(i', i) - \gamma = F_j(i', p') + g(i', i) - \gamma_j(i', i)$ by the induction hypothesis: it is one of the per-transcript DP's candidates for $(j, p)$. Conversely, every valid predecessor $(i', p')$ of $(j, p)$ lies in exactly one state of $i'$ by the induction hypothesis, remains after intersection with $i$'s color because $i$ has a placement in $j$, and lies in exactly one class of that state's refinement, yielding the same value. Thus the candidate sets coincide. Including the fresh-start score $w_i$ in both, the maximum for each pair is $F_j(i, p)$. Grouping pairs with equal maxima produces states that partition the pairs. □

**Proposition 4** (prefix sharing)**.** *If transcripts $j$ and $j'$ satisfy* $\mathrm{tpos}_{j'}(i) = \mathrm{tpos}_j(i) + c$ *for a constant $c$ and every anchor $i \le m$ (as multisets of placements), then $F_{j'}(i, p + c) = F_j(i, p)$ for all $i \le m$.*

*Proof.* Validity and cost depend on placements only through $\Delta t$, which is invariant under the shift, and anchors after $m$ are never predecessors of anchors up to $m$. The claim follows by induction on $i$. □

### S2 Additional results

#### S2.1 Mapping accuracy by read type

Table S1: Table 1 split by whether the simulated read reaches the transcript's 5′ end. Unique top-1: the true transcript is the only best-scoring hit; this cannot hold for reads from regions where isoforms are identical, so it is well below 100% for truncated reads. Other columns as in Table 1.

| Reads | Subset | Mapper | Top-1 | Unique top-1 | In set | Set size |
|---|---|---|---|---|---|---|
| ONT 95% | full length (0.95 M) | `spearfish` | 99.6% | 89.4% | 99.8% | 1.8 |
| | | `minimap2` | 99.7% | 89.4% | 99.9% | 3.9 |
| | 5′-truncated (1.02 M) | `spearfish` | 98.8% | 60.4% | 99.1% | 7.9 |
| | | `minimap2` | 98.7% | 60.5% | 99.1% | 10.4 |
| HiFi | full length (0.94 M) | `spearfish` | 99.9% | 89.6% | 100.0% | 2.4 |
| | | `spearfish`, ext. | 99.9% | 89.6% | 100.0% | 2.4 |
| | | `minimap2` | 99.7% | 89.6% | 99.7% | 3.0 |
| | 5′-truncated (1.01 M) | `spearfish` | 99.2% | 60.7% | 99.3% | 8.9 |
| | | `spearfish`, ext. | 99.3% | 60.8% | 99.4% | 8.9 |
| | | `minimap2` | 97.1% | 60.4% | 97.2% | 9.1 |
| ONT 90% | full length (0.11 M) | `spearfish` | 98.4% | 90.9% | 98.6% | 1.5 |
| | | `minimap2` | 98.4% | 90.9% | 98.7% | 3.5 |
| | 5′-truncated (0.12 M) | `spearfish` | 97.0% | 60.0% | 97.5% | 7.4 |
| | | `minimap2` | 96.9% | 60.0% | 97.4% | 10.3 |

#### S2.2 Mapping accuracy by transcript category

Because spike-in controls are synthetic and lack paralogs, pseudogenes, and long non-coding RNAs, in Table S2 we stratify the simulated reads of Table 1 by GENCODE transcript biotype and by cross-gene sequence sharing (measured as the largest fraction of a transcript's level-3 seed labels also present in any single other gene) to evaluate performance on multi-copy families. Reads from long non-coding, nonsense-mediated decay (NMD), and retained-intron transcripts are mapped as accurately as those from protein-coding transcripts. Wrong-gene assignments occur most often on pseudogenes and transcripts sharing at least 90% of their seeds with another gene (0.2–0.5% of reads for `spearfish` versus 0.1–0.4% for `minimap2`), though on HiFi reads `spearfish`'s top-1 accuracy stays above `minimap2`'s in every category.

Table S2: Table 1 by category of the source transcript. sf: `spearfish` (default preset; HiFi without span extension); mm2: `minimap2`; lrk: `lr-kallisto`. Upper rows: GENCODE v50 transcript type (pseudogene: all pseudogene types). Lower rows: largest fraction of the transcript's seed labels shared with one other gene. Wrong gene: the read is mapped but no best-scoring hit is in its gene. Top-1 and In set as in Table 1.

| Reads | Category | Reads | Top-1 | | | In set | Wrong gene | |
|---|---|---|---|---|---|---|---|---|
| | | | sf | mm2 | lrk | sf | sf | mm2 |
| ONT 95% | protein-coding | 862 k | 99.1% | 99.1% | 93.0% | 99.6% | 0.05% | 0.04% |
| | lncRNA | 581 k | 99.3% | 99.2% | 91.4% | 99.4% | 0.05% | 0.03% |
| | NMD | 278 k | 99.2% | 99.2% | 92.0% | 99.5% | 0.02% | 0.01% |
| | retained intron | 112 k | 99.4% | 99.4% | 92.7% | 99.5% | 0.07% | 0.06% |
| | pseudogene | 45 k | 99.0% | 99.0% | 90.9% | 99.1% | 0.20% | 0.15% |
| | shared < 10% | 1.37 M | 99.3% | 99.3% | 93.1% | 99.5% | 0.01% | 0.00% |
| | shared 10–50% | 266 k | 99.1% | 99.2% | 90.3% | 99.4% | 0.02% | 0.00% |
| | shared 50–90% | 145 k | 99.1% | 98.7% | 89.3% | 99.4% | 0.05% | 0.03% |
| | shared ≥ 90% | 185 k | 98.8% | 98.7% | 91.1% | 99.3% | 0.46% | 0.38% |
| HiFi | protein-coding | 850 k | 99.6% | 98.5% | 96.6% | 99.7% | 0.03% | 0.02% |
| | lncRNA | 571 k | 99.5% | 98.2% | 96.5% | 99.6% | 0.05% | 0.03% |
| | NMD | 274 k | 99.5% | 98.6% | 96.0% | 99.6% | 0.02% | 0.01% |
| | retained intron | 111 k | 99.6% | 98.4% | 97.2% | 99.7% | 0.06% | 0.05% |
| | pseudogene | 44 k | 99.4% | 95.8% | 98.4% | 99.5% | 0.16% | 0.12% |
| | shared < 10% | 1.35 M | 99.6% | 98.5% | 97.2% | 99.7% | 0.01% | 0.00% |
| | shared 10–50% | 262 k | 99.6% | 98.0% | 95.6% | 99.6% | 0.01% | 0.00% |
| | shared 50–90% | 143 k | 99.5% | 98.0% | 94.2% | 99.6% | 0.03% | 0.01% |
| | shared ≥ 90% | 182 k | 99.2% | 97.8% | 95.6% | 99.5% | 0.34% | 0.27% |
| ONT 90% | protein-coding | 103 k | 98.1% | 98.0% | 66.5% | 98.6% | 0.02% | 0.01% |
| | lncRNA | 74 k | 97.1% | 97.2% | 46.5% | 97.2% | 0.08% | 0.02% |
| | NMD | 34 k | 97.9% | 98.0% | 63.8% | 98.4% | 0.02% | 0.00% |
| | retained intron | 12 k | 97.9% | 97.8% | 58.0% | 98.0% | 0.02% | 0.00% |
| | pseudogene | 5 k | 96.7% | 96.8% | 44.0% | 96.8% | 0.19% | 0.09% |
| | shared < 10% | 175 k | 97.8% | 97.8% | 59.1% | 98.1% | 0.01% | 0.00% |
| | shared 10–50% | 31 k | 97.4% | 97.0% | 54.7% | 97.7% | 0.08% | 0.01% |
| | shared 50–90% | 19 k | 97.4% | 97.3% | 53.9% | 97.7% | 0.12% | 0.03% |
| | shared ≥ 90% | 14 k | 97.5% | 97.5% | 58.0% | 97.9% | 0.25% | 0.19% |

##### S2.3 Reads with library structure

PBSIM3 samples reads from transcript sequence alone. To test realistic untrimmed cDNA libraries, we added the ONT cDNA strand-switching and VN primers (or PacBio Iso-Seq primers) to both ends of 90% of ONT reads and 98% of HiFi reads, trimming each primer by 0–5 nt at its outer end. We also appended a 20–60 nt poly(A) tail to 84% of ONT reads and 95% of HiFi reads reaching their transcript's 3′ end (matching the fractions of SG-NEx cDNA and PacBio reads with a 15-base A- or T-run near an end), shifting the cleavage site 0–10 nt upstream of the annotated end. All added bases include substitutions at the platform's error rate (5% or 0.5%), and minus-strand reads receive the reverse complement (poly(T) at the start). Ground-truth transcript counts remain unchanged.

On these reads, `spearfish` + `oarfish` is more accurate than `minimap2` + `oarfish` on every quantification metric, by a slightly wider margin than on clean reads (Table S3, Table 3), where the 3′-end evidence comes from the end fit (Section 5.4) rather than from the poly(A) tail. Mapping accuracy shifts less (Table S4). On ONT 95% reads, `spearfish` ranks the true transcript first for 98.3% of reads versus 97.4% for `minimap2`, and both include the true transcript in their near-best set for 99.3% of reads. Top-1 accuracy drops slightly for both mappers relative to clean reads (by 0.9 and 1.7 percentage points), mainly because primer bases can match transcript sequence past an isoform's end by chance. Meanwhile, `spearfish` identifies the true transcript uniquely more often than on clean reads (76.6% versus 74.3%) because the poly(A) tail locates the 3′ end. At 90% accuracy, `spearfish` is ahead on top-1 by 0.7 percentage points, and both mappers include the true transcript in their near-best set for 97.9% of reads.

Table S3: Table 3 on the simulated reads with primers and poly(A) tails added.

| Reads | Pipeline | Spearman all / expr. | MARD all / expr. |
|---|---|---|---|
| ONT 95% | `spearfish` + `oarfish` | **0.911** / **0.874** | **0.115** / **0.158** |
| | `minimap2` + `oarfish` | 0.877 / 0.821 | 0.139 / 0.197 |
| | `lr-kallisto` | 0.779 / 0.715 | 0.251 / 0.366 |
| HiFi | `spearfish` + `oarfish` | **0.914** / **0.876** | **0.111** / **0.155** |
| | `minimap2` + `oarfish` | 0.881 / 0.824 | 0.135 / 0.194 |
| | `lr-kallisto` | 0.835 / 0.785 | 0.180 / 0.236 |

Table S4: Table 1 on the simulated reads with primers and poly(A) tails added. Columns as in Table 1.

| Reads | Mapper | Mapped | Top-1 | In set | Set size | Gene |
|---|---|---|---|---|---|---|
| ONT 95% | `spearfish` | **99.5%** | **98.3%** | **99.3%** | **4.8** | **99.3%** |
| | `minimap2` | 99.4% | 97.4% | **99.3%** | 7.3 | 99.1% |
| | `lr-kallisto` | 89.8% | n/a | 88.6% | 7.1 | 89.7% |
| HiFi | `spearfish` | 99.2% | 98.7% | 99.2% | 5.5 | 99.1% |
| | `spearfish`, ext. | **99.7%** | **99.1%** | **99.6%** | 5.6 | **99.5%** |
| | `minimap2` | 98.4% | 97.5% | 98.3% | 6.2 | 98.2% |
| | `lr-kallisto` | 91.6% | n/a | 90.8% | **4.5** | 91.4% |
| ONT 90% | `spearfish` | **98.1%** | **97.0%** | **97.9%** | **4.4** | **98.0%** |
| | `minimap2` | 98.0% | 96.3% | **97.9%** | 7.1 | **98.0%** |
| | `lr-kallisto` | 58.4% | n/a | 57.7% | 20.1 | 58.3% |

##### S2.4 Agreement with `minimap2` on experimental reads

Table S5: `spearfish` (sf), `minimap2` (mm2) and `lr-kallisto` (lrk) on 200,000 random reads per SG-NEx run. Mapped: best hit spans at least half the read (`lr-kallisto`: the read pseudoaligns); on direct cDNA, many reads are below half for the aligners because of adapter and chimeric sequence. Jaccard index and set size: over reads mapped by both compared tools, using each tool's near-best set (hits within 95% of its best score; `lr-kallisto`: its equivalence class). Over the same reads, the `spearfish` and `minimap2` sets share a transcript for 99.0–99.7% of reads, and the `lr-kallisto` and `minimap2` sets for 96.7–99.8%; a `spearfish` best hit lies in the gene of `minimap2`'s primary alignment for 98.3–99.6%.

| | Mapped | | | Jaccard with mm2 | | Set size | | |
|---|---|---|---|---|---|---|---|---|
| Run | sf | mm2 | lrk | sf | lrk | sf | mm2 | lrk |
| H9 direct RNA | 97.6% | 98.0% | 66.8% | 0.57 | 0.54 | 7.1 | 13.5 | 21.3 |
| HEYA8 direct RNA | 98.3% | 98.7% | 67.7% | 0.55 | 0.53 | 7.5 | 14.3 | 22.2 |
| HEYA8 direct cDNA | 76.2% | 62.9% | 65.2% | 0.61 | 0.55 | 8.7 | 14.4 | 17.3 |
| HEYA8 PCR-cDNA | 93.5% | 91.9% | 67.5% | 0.53 | 0.46 | 6.2 | 12.5 | 16.5 |
| H9 PCR-cDNA | 89.5% | 86.5% | 65.6% | 0.56 | 0.49 | 6.4 | 12.4 | 16.1 |
| HEYA8 PacBio | 94.0% | 94.9% | 75.9% | 0.82 | 0.34 | 9.0 | 11.4 | 2.2 |
| K562 PacBio | 93.0% | 93.5% | 75.3% | 0.85 | 0.49 | 7.3 | 9.3 | 2.0 |

##### S2.5 Ablations

In Table S6, we disable one component of `spearfish` at a time on the simulated ONT 90% dataset, the hardest read set. Base-level tie-breaking has the largest impact on quantification: disabling it lowers top-1 accuracy by 4.8 percentage points and lowers expressed-transcript Spearman correlation from 0.847 to 0.787, while saving 14% of instructions. Hierarchical rescue has the largest effect on mapping sensitivity (top-1 falls from 97.7% to 91.6% without it) and accounts for 36% of mapping instructions. Its level-2 re-chaining, global triple table, and antisense rescue contribute 0.7, 0.9, and 0.1 percentage points of top-1, respectively.

Among scoring and alignment choices, low-score rejection reduces top-1 by 0.2 percentage points while raising overall Spearman correlation from 0.914 to 0.922 without affecting expressed transcripts, indicating that rejected reads are mainly false placements onto unexpressed transcripts. Aligning read ends and internal segments in uncompressed rather than HPC sequence improves every metric, and capping the end bonus at the shortest candidate adds 0.1 points of top-1. Replacing affine-gap alignment with unit-cost banded bit-parallel edit distance [46, 47] (or scoring orientation by edit distance) stays within 0.003 of the default estimates. Forcing read ends without soft-clipping matches clipped accuracy on simulated reads (which lack adapters) at 4% higher instruction count, whereas on experimental direct cDNA soft-clipping raised Spearman correlation from 0.78 to 0.84 (Section 5.3).

Indexing and chaining optimizations primarily affect speed. Colinear anchor merging and the choice of chainer affect only runtime cost: disabling colinear merging increases instructions by 12%, and switching from the prefix chainer to the refinement DP increases them by 91%. Capping LCP's run rule at 4 (Section S1.1) bounds extended windows in bases for 2% more instructions and no measurable loss in accuracy. Minimizer seeds match LCP within 0.1 percentage points on top-1 and 0.002 on quantification for 5% more instructions, whereas collapsing seed labels globally doubles the instruction count for no accuracy gain. Extending reported spans to read ends adds 1% to instructions and leaves simulated estimates unchanged (top-1 dips slightly without extension only because hits spanning under half the read count as unmapped), though span extension matters for `oarfish`'s span filters on experimental cDNA (Section 6.3).

Table S6: Ablations on 239,000 simulated ONT reads at 90% mean accuracy, one component changed per row and all rows quantified by `spearfish`'s standalone EM, which weights hits by their mapping score. Instructions: user-space instructions retired by mapping, which do not depend on machine load. The state-DP chainer's output differs from the default's in 53 of 3.4 M lines. †Measured with a build of `spearfish` 0.3.1 with the cap; with its default parse, mapping is identical and takes 528 G, the base of the +2%.

| Variant | Top-1 | Spearman all / expr. | MARD all / expr. | Instructions | Change |
|---|---|---|---|---|---|
| `spearfish` (default) | 97.7% | 0.922 / 0.847 | 0.0165 / 0.191 | 524 G | |
| no tie-break | 92.9% | 0.884 / 0.787 | 0.0233 / 0.252 | 449 G | −14% |
| tie-break on whole reads | 96.7% | 0.900 / 0.815 | 0.0205 / 0.228 | 827 G | +58% |
| unit-cost tie-break | 97.7% | 0.922 / 0.848 | 0.0165 / 0.188 | 520 G | −1% |
| read ends forced (no clipping) | 97.5% | 0.922 / 0.846 | 0.0164 / 0.190 | 543 G | +4% |
| end bonus not cut | 97.5% | 0.922 / 0.846 | 0.0165 / 0.191 | 524 G | 0% |
| read ends in HPC space | 96.8% | 0.917 / 0.841 | 0.0173 / 0.197 | 487 G | −7% |
| read middles in HPC space | 97.2% | 0.919 / 0.844 | 0.0169 / 0.194 | 496 G | −5% |
| orientation by edit distance | 97.7% | 0.922 / 0.847 | 0.0165 / 0.191 | 528 G | +1% |
| spans not extended to read ends | 97.5% | 0.922 / 0.847 | 0.0165 / 0.191 | 518 G | −1% |
| no low-score rejection | 97.9% | 0.914 / 0.847 | 0.0176 / 0.191 | 539 G | +3% |
| no rescue | 91.6% | 0.901 / 0.810 | 0.0213 / 0.260 | 336 G | −36% |
| rescue without re-chaining | 97.0% | 0.919 / 0.843 | 0.0170 / 0.197 | 391 G | −25% |
| rescue without the global table | 96.8% | 0.920 / 0.845 | 0.0169 / 0.195 | 447 G | −15% |
| rescue without antisense transcripts | 97.5% | 0.921 / 0.846 | 0.0166 / 0.192 | 508 G | −3% |
| no colinear merging | 97.7% | 0.922 / 0.847 | 0.0165 / 0.191 | 589 G | +12% |
| state-DP chainer (Theorem 3) | 97.7% | 0.922 / 0.847 | 0.0165 / 0.191 | 1,000 G | +91% |
| LCP with runs capped at 4 (Section S1.1)† | 97.7% | 0.922 / 0.847 | 0.0165 / 0.191 | 539 G | +2% |
| minimizer seeds ($k = 19, w = 23$) | 97.7% | 0.921 / 0.847 | 0.0166 / 0.191 | 548 G | +5% |
| global label collapsing | 97.6% | 0.921 / 0.847 | 0.0166 / 0.192 | 1,064 G | +103% |

##### S2.6 `lr-kallisto`'s sequin estimates on PacBio

Inspecting the sequin quantification results shows that `lr-kallisto`'s lower sequin MARD on the two PacBio runs is due to reads it discards. Under every pipeline, including `lr-kallisto`, both PacBio libraries exhibit strong molecule-length bias relative to the nominal spike-in mix table: sequins under 700 nt receive 2–3% of expected reads, 700–1,000 nt sequins 7–11%, 1–1.7 kb sequins 53–66%, and sequins over 1.7 kb 4.4–4.7 times as many reads as expected. Most of `lr-kallisto`'s apparent MARD advantage comes from four sequin genes for which it retains fewer than half the reads assigned by `spearfish`.

The largest contributor is gene R1_33. On K562, its 2.4 kb isoform R1_33_1 has roughly 19 times the reads expected from the mix table (an estimated 8,929 versus 480), while its equimolar 1.0 kb sibling R1_33_2 has about a third (143). The R1_33_1 reads are full length, highly accurate, and compatible only with R1_33_1 because they contain its 3′ exons, which are absent from R1_33_2. However, they also include two bases upstream of R1_33_1′s annotated 5′ end that match R1_33_2, whose first exon begins further upstream. Consequently, each read contains two 63-mers unique to R1_33_2, so intersecting all $k$-mer transcript sets yields an empty intersection. `lr-kallisto` then falls back to modal recovery (selecting the most frequent equivalence class among $k$-mer hits). As implemented, however, modal recovery tallies a class only when a left-to-right scan along the read transitions to a different class. Because the dominant R1_33_1 class runs all the way to the end of the read, the fallback returns nothing and the read is discarded. Of

8,991 reads mapping to R1__33__1, `lr-kallisto` discards 80% and retains only 1,789 (mostly 5′-truncated reads without those upstream bases). Trimming the first 55 bases restores pseudoalignment for 97% of the discarded reads.

Discarding those reads lowers MARD only because R1__33__1 is strongly over-represented in the library relative to the mix table. Excluding these four genes halves the sequin MARD gap and shrinks the Spearman gap from 0.020 to 0.010 on K562 and from 0.012 to 0.002 on HEYA8. A related effect explains `lr-kallisto`'s flatter SIRV-4 profile on H9 PCR-cDNA: it retains markedly fewer reads than `spearfish` for short isoforms (under 600 nt, which are over-represented after PCR) and slightly more for isoforms over 1 kb, again discarding reads because of the same isoform-end offsets that `spearfish` handles via clipped, uncompressed read ends (Section 5.3).

#### S2.7 CPU time and memory

Table S7 reports total CPU time and peak resident memory (RSS) for the runs in Table 2.

Table S7: CPU seconds and peak resident memory. `oarfish`'s memory is dominated by its per-read alignment store, which it keeps for per-read output even when the EM runs on equivalence classes.

| Run | `spearfish` map CPU s | `minimap2` CPU s | `spearfish` RSS | `minimap2` RSS | `spearfish` + `oarfish` RSS | `lr-kallisto` RSS |
|---|---|---|---|---|---|---|
| H9 direct RNA | 50 | 2,882 | 3.2 GB | 19.0 GB | 8.7 GB | 4.2 GB |
| HEYA8 direct RNA | 85 | 5,050 | 3.2 GB | 20.2 GB | 8.5 GB | 4.2 GB |
| HEYA8 direct cDNA | 1,870 | 69,927 | 3.9 GB | 19.4 GB | 8.3 GB | 4.2 GB |
| HEYA8 PCR-cDNA | 3,632 | 111,583 | 3.7 GB | 20.2 GB | 9.3 GB | 4.2 GB |
| H9 PCR-cDNA | 4,173 | 102,193 | 3.8 GB | 19.3 GB | 8.2 GB | 4.2 GB |
| HEYA8 PacBio | 406 | 41,916 | 1.8 GB | 13.7 GB | 8.2 GB | 4.2 GB |
| K562 PacBio | 223 | 20,469 | 1.8 GB | 13.6 GB | 8.4 GB | 4.2 GB |
| Simulated ONT | 183 | 13,830 | 3.5 GB | 16.6 GB | 8.3 GB | 4.2 GB |
| Simulated HiFi | 126 | 10,453 | 1.9 GB | 13.5 GB | 8.4 GB | 4.2 GB |

#### S2.8 Allocation of mapping time

Table S8 breaks down `spearfish map` CPU time by stage across five experimental runs using call-stack profiling (`perf`, 499 Hz, 32 threads). Tie-breaking accounts for 21–27% of mapping time. Ties without a globally shared anchor (paralogs and repeats, handled by anchor-sharing groups in Section 5.3) take 2–7% on ONT reads and 2–3% on PacBio reads. Across tie-breaking and orientation checks, alignment kernels consume 11–21% of total CPU time. Hierarchical rescue accounts for 32–44% on ONT cDNA and 27% on direct RNA as it re-seeds low-coverage or uncertain reads at level 2.

Table S8: Share of mapping CPU time by stage (`spearfish`'s default preset for each run, 32 threads; samples attributed by call stack). Rescue is off in the HiFi preset, and span extension in the HiFi and direct RNA presets. The profiled build is the evaluated source compiled with frame pointers and with the tie-break kept out of line, so its stages can be told apart; neither changes the output.

| **Stage** | **K562 PacBio** | **HEYA8 PacBio** | **HEYA8 direct cDNA** | **H9 PCR-cDNA** | **HEYA8 direct RNA** |
|---|---|---|---|---|---|
| Seeding (LCP parse) | 23.1% | 21.8% | 6.0% | 6.8% | 9.0% |
| Anchor selection | 20.9% | 20.9% | 5.7% | 7.1% | 8.4% |
| Chaining | 24.9% | 25.1% | 9.2% | 12.6% | 18.4% |
| Rescue | 0.0% | 0.0% | 43.7% | 31.7% | 27.4% |
| Tie-break: shared anchors | 17.3% | 18.9% | 17.2% | 20.0% | 23.7% |
| Tie-break: no shared anchor | 2.7% | 2.4% | 2.7% | 6.5% | 1.6% |
| Tie-break: other | 1.1% | 1.2% | 0.6% | 0.9% | 1.4% |
| Orientation | 0.2% | 0.2% | 1.7% | 0.8% | 0.1% |
| Span extension | 0.0% | 0.0% | 2.7% | 2.6% | 0.0% |
| Other mapping | 6.7% | 6.4% | 8.9% | 9.1% | 5.9% |
| Output records | 2.6% | 2.5% | 1.5% | 1.7% | 3.3% |
| Input, threads, setup | 0.6% | 0.5% | 0.1% | 0.1% | 0.9% |
| Tie-break, total | 21.1% | 22.6% | 20.5% | 27.4% | 26.7% |
| Alignment kernels | 10.7% | 11.4% | 16.4% | 20.7% | 14.3% |

##### S2.9 Mapping time at 16 threads

At 16 threads, `spearfish` maps 24–100× faster than `minimap2` and 2.5–10.3× faster than `lr-kallisto`'s pseudoalignment step (Table S9), matching the 32-thread ratios (Table 2). Doubling threads from 16 to 32 nearly doubles speed for all three tools: 1.92–1.98× for `spearfish`, 1.78–1.95× for `minimap2`, and 1.63–1.97 × for `kallisto bus` (lowest on short runs, where loading its index takes about 5 s).

Table S9: Mapping wall-clock seconds at 16 threads, as in Table 2: sf, `spearfish map`; mm2, `minimap2` (index prebuilt, output compressed to BAM); lrk, `kallisto bus`. The last three columns give each mapper's speedup from 16 to 32 threads: its time at 16 threads divided by its time at 32 (Table 2).

| | **16 threads** | | | | | **Speedup, 16 → 32 threads** | | |
|---|---|---|---|---|---|---|---|---|
| Run | sf | mm2 | lrk | mm2 ÷ sf | lrk ÷ sf | sf | mm2 | lrk |
| H9 direct RNA | 3.1 | 182 | 32 | 59× | 10.3× | 1.92× | 1.82× | 1.73× |
| HEYA8 direct RNA | 5.3 | 311 | 49 | 59× | 9.3× | 1.93× | 1.86× | 1.79× |
| HEYA8 direct cDNA | 116 | 3,836 | 285 | 33× | 2.5× | 1.97× | 1.78× | 1.93× |
| HEYA8 PCR-cDNA | 226 | 6,625 | 780 | 29× | 3.5× | 1.98× | 1.95× | 1.97× |
| H9 PCR-cDNA | 259 | 6,086 | 737 | 24× | 2.8× | 1.98× | 1.95× | 1.96× |
| HEYA8 PacBio | 25 | 2,523 | 110 | 100× | 4.4× | 1.96× | 1.93× | 1.88× |
| K562 PacBio | 13.8 | 1,239 | 69 | 90× | 5.0× | 1.96× | 1.94× | 1.86× |
| Simulated ONT | 11.4 | 843 | 64 | 74× | 5.6× | 1.95× | 1.93× | 1.78× |
| Simulated HiFi | 7.8 | 637 | 30 | 81× | 3.8× | 1.96× | 1.94× | 1.63× |

##### S2.10 A truncation model for `oarfish`

Because estimated abundances on PCR-cDNA fall with isoform length relative to spike-in concentrations (Section 6.3), we added a truncation model (`--model-truncation`) to `oarfish`. Under this model, each read's placement on a transcript is drawn from a four-component mixture of 3′-anchored, 5′-anchored, full-length, and uniformly placed reads. The EM fits the mixture weights, read-length distributions, and gap distributions between read and transcript ends (modeled nonparametrically on log-spaced bins with a component scaling with transcript length), so no protocol-specific setting is needed. Incorporating this placement probability into each alignment's likelihood assigns a read that is full length for a short isoform, yet truncated for a longer one, preferentially to the shorter isoform to the extent that such truncation is unlikely under the fitted distributions. Because read-end coordinates are needed for this model, `oarfish` enables span extension for `spearfish` across all presets whenever `--model-truncation` is active.

With `spearfish` mappings, the truncation model lowers SIRV MARD on all seven SG-NEx runs (from 0.145 to 0.125 on H9 direct RNA, from 0.422 to 0.418 on HEYA8 PCR-cDNA, and by 0.006–0.008 on the two PacBio runs) while changing sequin MARD by at most 0.005. On H9 PCR-cDNA, however, the spread across SIRV-4 isoforms is unchanged (coefficient of variation 1.18 with and without the model), indicating that PCR amplification bias remains the primary driver there.

##### S2.11 Accuracy of fold-change estimation in different transcript categories

In Figure S1, we compare the accuracy of estimated expression fold changes derived from `spearfish`, `minimap2`, and `lr-kallisto` mappings across transcript categories that are difficult for read assignment. The simulated two-condition libraries contain 360 transcripts spanning paralogs, processed pseudogenes and their parental transcripts, lncRNAs, NMD isoforms and coding partners, retained-intron isoforms and coding partners, and nested coding transcripts and hosts, generated as ONT cDNA reads (95% mean accuracy) and Iso-Seq HiFi reads. To isolate mapper accuracy, candidate mappings from all three tools are quantified with the same EM estimator (`spearfish em` from v0.3.2, reading the quantification-score column), which weights `spearfish` mappings by their quantification scores (Section 5.4) as `oarfish` does and weights `minimap2` and `lr-kallisto` mappings by their scores. We summarize errors across 140 gene or paralog-pair units per tool and platform, reporting absolute error in $\log_2$ fold change for individual targets and root-mean-square error across each paralog pair, with $\log_2$ fold changes computed as $\log_2\left(\frac{B+1}{A+1}\right)$ from condition counts $A$ and $B$ scaled to each library's true read total.

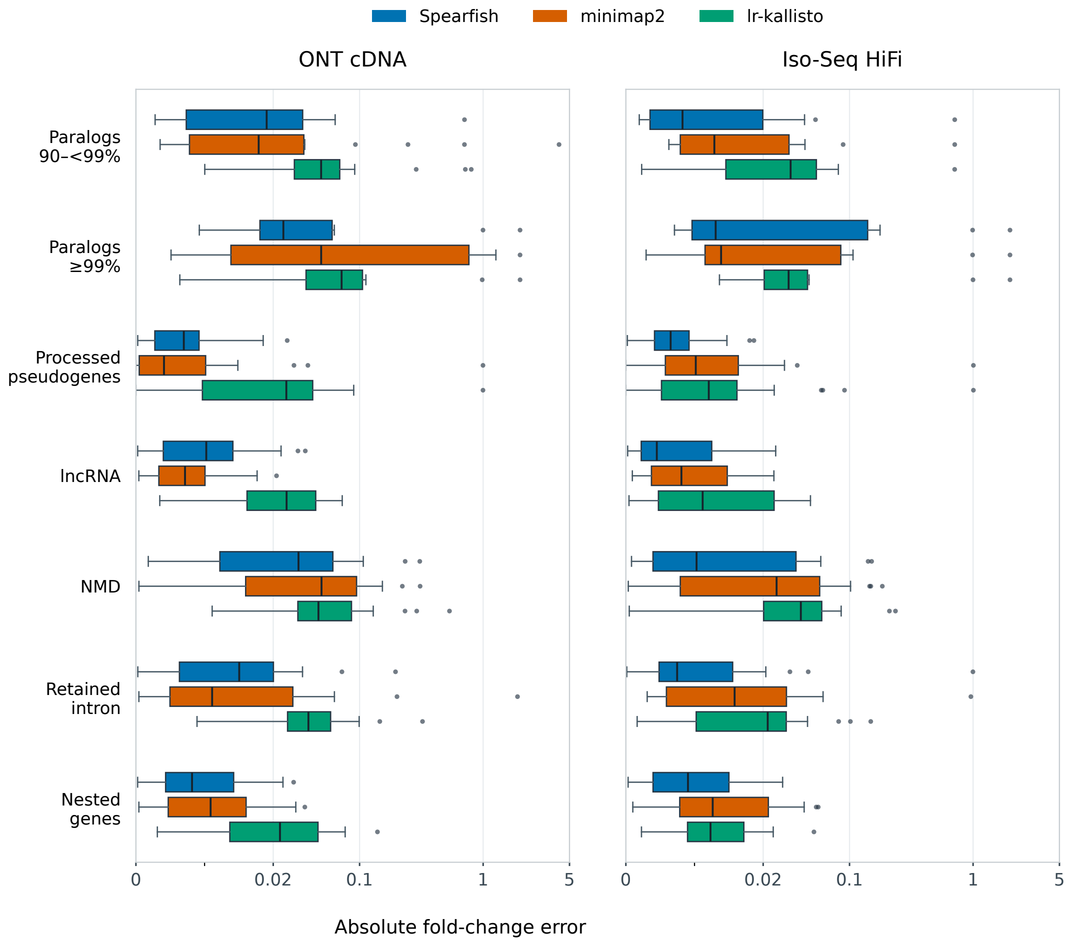


Figure S1: Error in estimated $\log_2$ expression fold change by transcript category, on simulated ONT cDNA reads (95% mean accuracy) and Iso-Seq HiFi reads. Mappings from all three tools are quantified with the same `spearfish` EM estimator (`spearfish`'s mappings weighted by their quantification scores). Errors are absolute errors for individual targets and root-mean-square errors across paralog pairs (lower is better). Boxes show the interquartile range, with the median marked inside. Points beyond the whiskers show outliers.

#### S3 Default parameters and heuristic rules

Table S10 lists the numeric thresholds, scoring bonuses, and preset-specific rules omitted from the main text of Section 5, together with the motivation for each. Chaining parameters from Section 5.1 and EM settings from Section 5.4 are included so that all defaults appear in one place.

Table S10: Default parameters, heuristic thresholds, and preset-specific rules in `spearfish` 0.3.1.

| Stage | Rule | Default / threshold | Rationale |
|---|---|---|---|
| **Gene filtering** | Repetitive seed cap | Ignore level-3 labels in $> 1000$ genes | Prevents ubiquitous low-complexity or repeat seeds from flooding gene scoring. |
| | Score floor | Drop genes $< 50\%$ of top greedy score | Eliminates incidental seed matches far below the best-supported locus. |
| | Rank & tie rule | Top 8 genes plus ties within 1% of 8th (up to 64) | Avoids arbitrary cutoffs in multi-copy families (e.g. subtelomeric genes) with identical scores. |
| | Strand mode | Both strands; sense only in `drna` | Direct RNA reads are strand-specific; cDNA protocols sequence both strands. |
| **Chaining** (Section 5.1) | Anchor weight $w_i$ | Span capped at 100; slightly lowered for multi-gene seeds | Caps influence of long merged runs and weights gene-specific seeds higher. |
| | Gap bounds & cost | $\alpha = 1.5$, $h = 50$, $o_{\max} = 100$, $g_{\max} = 5000$; no log gap term | Linear penalty on $\lvert\Delta q - \Delta t_j\rvert$ distinguishes isoforms; introns are absent in transcripts. |
| | Beam width | None (exact DP) | Pruning low-scoring states would drop near-best candidates used by EM. |
| **Level-2 rescue** (Section 5.2) | Preset availability | Enabled in `ont` and `drna`; off in `hifi` | HiFi reads ($\approx$0.1% error) rarely lose level-3 seeds, so rescue adds no sensitivity. |
| | Low coverage | Best chain $< 50\%$ of compressed read $q$; also rescue $H'$ if within 30 pt or above min score | Targets noisy reads whose level-3 cores are disrupted by sequencing errors. |
| | Global triples | If still $< 50\%$ covered, or rescued within 10 pt of min score: runs of 3 level-2 seeds | Triples are specific enough to index globally when level-3 seeding misses the true gene. |
| | Uncertain re-chain | Rescued, $< 3$ anchors, $< 30\%$ anchored, or $H'$ within 65% of $H$: up to 256 $H$ and 16 antisense $H'$ hits | Refines sparse or ambiguous chains at level 2 without re-chaining confident hits. |
| **Tie-break & ends** (Section 5.3) | Tie-break set $T$ | Hits within 5% (or 30 pt) of $s^*$, up to 256; skip if $\lvert T\rvert < 2$ | Restricts base-level alignment to plausible near-ties that can affect EM weights. |
| | Extension & clipping | `minimap2` `map-ont`/`map-hifi` affine scores; X-drop 50; end bonus 5 (capped at shortest candidate); fixed diagonal end bands on `hifi` | Soft-clips untrimmed adapters/primers and prevents stray flanks from inflating longer isoforms' scores. |
| | Uncompressed seq. | Read ends uncompressed except `drna`; internal segments also uncompressed on `ont` | Prevents template-switch `G`s (absent in direct RNA) from merging with a leading `G` in HPC space. |
| | Poly(A) detection | Past last anchor: 15-bp window with $\leq 1$ non-A within 80 bp of read end (10 bp in `drna`) | Stops alignment before poly(A) tails; `drna` lacks long cDNA primers past the tail. |
| | Poly(A) 3′ bonus | +10 pt for candidates ending within $15 + 0.1 \times$ (read end) bp of tail start, or with an A-run there | Adds weight to isoforms whose annotated 3′ cleavage site matches the detected tail boundary. |
| | Orientation | Triggered if $H, H'$ best chains within 35% (or 30 pt) or identical opposite-strand hits; clipped score +30 pt for tail ($\leq 80$ bp flank); `ont` local align if lead $\leq 150$ pt | Resolves sense/antisense overlaps while keeping both orientations; local alignment breaks close cDNA margins. |
| | Quant. orientation | Identical opposite-strand pairs: exact local align; tail at 3′ end credited with $\leq 120$ bp flank | Handles nested antisense isoforms and longer untrimmed adapter flanks for EM without altering mapping scores. |
| | HiFi tail shortcut | If one orientation has a poly(A) tail, map it first; check other only if hit spans $<$ 80% of read | Avoids unnecessary reverse-complement work on accurate reads with clear strand evidence. |
| **Output & EM** (Section 5.4) | Minimum score | Unmapped if best chain $< 36 + 4\ln(\ell/1000)$; short/borderline rescued hit also requires clipped score $\geq 40 +$ (read span) | Scales logarithmically with read length $\ell$ because longer reads have more chances to form spurious short chains. |
| | Reporting window | All transcripts within 30% (or 30 pt) of $s^*$; exact ties ordered by read/transcript hash | Retains every candidate needed downstream and avoids systematic index-order tie bias. |
| | End fit (EM only) | +5 pt to quantification score when untailed, right end reaches farthest and within 4 compressed bases of 3′ end (when 3′ ends differ by $> 4$) | Uses flush 3′-end alignment as isoform evidence in untailed reads without altering mapping scores. |
| | `oarfish` EM | $P(r \mid t_j) \propto \exp\big((s_j - s_{\max}(r))/D\big)$ over hits $\geq 95\%$ of $s_{\max}(r)$, $D = 2$ | Default `oarfish` likelihood scaling shared across compared pipelines. |

#### S4 Pseudocode

Algorithms S1–S3 summarize single-read mapping (Section 5), prefix chaining (Section 5.1), and base-level tie-breaking (Section 5.3). Steps appear in implementation order. Caches, SIMD batching, and interval-reporting bookkeeping are omitted, and numeric thresholds follow Table S10.

Algorithm S1: MapRead: map a single read $r$ against index $I$ under preset $P$.

**Input:** read $r$, core-graph index $I$, preset $P \in \{\texttt{ont}, \texttt{drna}, \texttt{hifi}\}$
**Output:** set $H$ of scored candidate mappings (with mapping and quantification scores), or $\emptyset$

**for** orientation $o \in$ Orientations$(r, P)$ **do** ▷ sense only in drna; poly(A) first in hifi
  $q_o \leftarrow$ Hpc$(r_o)$; $S_o \leftarrow$ LcpParse$(q_o, 3)$ ▷ level-3 cores
  $A \leftarrow$ LookupSeeds$(S_o, I)$ ▷ skip labels in $> 1000$ genes
  $G \leftarrow$ FilterGenes$(A)$ ▷ greedy coverage; top 8 + ties, Table S10
  $H_o \leftarrow \emptyset$
  **for** each gene $g \in G$ **do**
    $A_g \leftarrow$ MergeColinear$(A[g])$ ▷ merge runs within one unitig
    $H_o \leftarrow H_o \cup$ PrefixChain$(A_g)$ ▷ Algorithm S2; $F_j$ for all $j \in g$
  **if** $P = \texttt{hifi}$ **and** single-tail shortcut satisfied **then break** ▷ span $\geq 80\%$; Table S10
$(H, H') \leftarrow$ OrderByBestScore$(H_+, H_-)$
**if** $P \neq \texttt{hifi}$ **and** cov$(H) < 0.5\ |q|$ **then** ▷ level-2 rescue (Section 5.2)
  $(H, H') \leftarrow$ RescueLocal$(q, G, H, H')$ ▷ candidate-gene level-2 seeds
  **if** cov$(H) < 0.5\ |q|$ **or** borderline rescue **then**
    $(H, H') \leftarrow$ RescueTriples$(q, I, H, H')$ ▷ global level-2 triples
**if** $P \neq \texttt{hifi}$ **and** ChainUncertain$(H, H')$ **then** ▷ Table S10
  $(H, H') \leftarrow$ RechainLevel2$(q, H, H')$ ▷ $\leq 256$ $H$ hits, $\leq 16$ antisense $H'$ hits
$H \leftarrow$ TieBreak$(r, H)$ ▷ Algorithm S3
$\delta_{\text{quant}} \leftarrow 0$
**if** AmbiguousStrand$(H, H')$ **then** ▷ within 35% / 30 pt or identical seq.
  $H' \leftarrow$ TieBreak$(r, H')$
  $(\Delta_{\text{strand}}, \delta_{\text{quant}}) \leftarrow$ CompareStrands$(r, H, H', P)$ ▷ clipped/local + poly(A); Table S10
  $H \leftarrow H \cup$ ShiftScores$(H', \Delta_{\text{strand}})$
$s^* \leftarrow \max_{h \in H} h.\text{score}$
**if** $s^* < \tau(|r|)$ **or** FailsRescueCheck$(H)$ **then return** $\emptyset$ ▷ min score; Table S10
$H \leftarrow \{h \in H : h.\text{score} \geq \max(0.7s^*, s^* - 30)\}$ ▷ reporting window
**for** each hit $h \in H$ **do** $h.\text{qscore} \leftarrow h.\text{score} + \delta_{\text{quant}}(h) +$ EndFit$(r, h, H)$ ▷ Table S10
**return** $H$

Algorithm S2: PrefixChain: exact chain scores for every candidate transcript of one gene (Proposition 4).

**Input:** merged anchors $A_g = (a_1, \dots, a_n)$ of gene $g$, ordered by query start
**Output:** exact colinear chaining optimum $F_j$ for every candidate transcript $j$ in $g$

**for** each transcript $j$ with a placement in $A_g$ **do**
  $P_j \leftarrow$ list of $(i, \text{tpos}_j(i))$ in anchor order (Eq. 1), shifted so its first placement is 0
sort the lists $P_j$ lexicographically; $P_{\text{prev}} \leftarrow \emptyset$
**for** each $P_j$ in sorted order **do**
  **if** $P_j = P_{\text{prev}}$ **then** $F_j \leftarrow F_{\text{prev}}$; **continue** ▷ identical shifted placements
  $c \leftarrow$ CommonPrefixLen$(P_j, P_{\text{prev}})$ ▷ reuse DP values $f[1..c]$
  **for** $k = c+1, \dots, |P_j|$ **do** ▷ entry $k$ is anchor $i_k$ at placement $p_k$
    $V \leftarrow \{k' < k : i_k - i_{k'} \leq h, \Delta q, \Delta t_j \in [-o_{\max}, g_{\max}]\}$
    $f[k] \leftarrow \max\big(w_{i_k}, \max_{k' \in V} f[k'] + \text{gain}(k', k) - \gamma_j(i_{k'}, i_k)\big)$ ▷ Eq. 3
  $F_j \leftarrow \max_k f[k]$; $P_{\text{prev}} \leftarrow P_j$
**return** $\{F_j\}$

Algorithm S3: TieBreak: re-rank near-best hits $H$ of one orientation by base-level alignment.

**Input:** read $r$, scored chains $H$ for one orientation
**Output:** re-ranked hits $H$

1 $s^* \leftarrow \max_{h \in H} h.\text{score}$
2 $T \leftarrow$ TopK($\{h \in H : h.\text{score} \geq \max(0.95s^*, s^* - 30)\}, 256$)
3 **if** $|T| < 2$ **then return** $H$
4 $A_{\text{shared}} \leftarrow \bigcap_{j \in T}$ ChainAnchors($j$)
5 **if** $A_{\text{shared}} \neq \emptyset$ **then**
6     $\text{cost}[j] \leftarrow 0$ for all $j \in T$
7     **for** each consecutive pair $(a, b)$ in $A_{\text{shared}}$ **do**
8         $\mathcal{G} \leftarrow$ PartitionBySeq($T, a, b$) ▷ group by transcript seq.
9         **if** $|\mathcal{G}| > 1$ **then** ▷ skip identical intervals
10             **for** each group $g \in \mathcal{G}$ **do**
11                 $c \leftarrow$ BandedGlobalAlign($r[a..b], \text{seq}(g)$) ▷ affine gap; Table S10
12                 **for** each $j \in g$ **do** $\text{cost}[j] \leftarrow \text{cost}[j] + c$
13     $p_{\text{tail}} \leftarrow$ DetectPolyA($r, A_{\text{shared}}$) ▷ Table S10
14     **for** each end $e \in \{\text{left}, \text{right}\}$ **do**
15         $\text{cost} \leftarrow \text{cost} +$ ExtendEnd($r, T, A_{\text{shared}}, e, p_{\text{tail}}$) ▷ clipped; raw seq. except `drna`
16 **else** ▷ paralogs, repeats: no global anchor
17     $\mathcal{G} \leftarrow$ GroupBySharedAnchors($T$)
18     run lines 6–15 independently within each group $g \in \mathcal{G}$
19     adjust inter-group $\text{cost}[j]$ by clipped chain-representative alignment
20 $c^* \leftarrow \min_{j \in T} \text{cost}[j]$
21 **for** each $j \in T$ **do** $s_j \leftarrow s^* - (\text{cost}[j] - c^*)$ ▷ score points
22 **return** SortByScore($H$)
